# Glial Mutant Huntingtin Initiates a Progressive Pathological Cascade Toward Neuronal Involvement in Drosophila

**DOI:** 10.64898/2026.10.21.760655

**Authors:** Debalina Chatterjee, Jassika Gupta, Mansi Yadav, Parul Mittal, Namita Agrawal, Anju Shrivastava

**Affiliations:** Department of Zoology, University of Delhi, North Campus, Chhatra Marg, Delhi-110007, India

**Author notes:** Corresponding Authors Prof. Namita Agrawal Professor Department of Zoology University of Delhi North Campus, Delhi-110007, India Prof. Anju Shrivastava Professor Department of Zoology University of Delhi North Campus, Delhi-110007, India.

**Keywords:** Huntington’s disease, Glia, *Drosophila melanogaster*, glia-neuron interaction, protein aggregation

## Abstract

The mechanisms of involvement of glial cells in the progression of Huntington’s disease remain sparingly elucidated. Using the *Drosophila melanogaster* model, we investigated the temporal course of the disease pathogenesis induced by pan-glial expression of HTTex1pQ93 in comparison to the temporal course of pathology induced by pan-neuronal expression. Our behavioural data were supported by quantitative confocal microscopy, quantification of aggregate formation and its morphometry, Mander’s M2 colocalization index, and correlative holotomography. Expression of mHTT in glial cells led to developmental defects and subsequent delayed onset of motor dysfunction and death with increasing amounts of HTT and changes in the immunoreactivity of glial cells. In spite of the progressive increase in HTT accumulation, glial aggregates showed stability after their initial formation, while neuronal mHTT revealed distinct age-dependent remodelling of its aggregates. Importantly, there was an initial preference of glial mHTT for glial compartments followed by a progressively increasing number of HTT in the neurons with aging. Correlative holotomography provided spatial evidence of the neuronal association of HTT.

Together, these findings demonstrate that glial mHTT establishes a distinct progressive pathological trajectory characterised by evolving glial pathology, increasing neuronal involvement, and delayed functional decline, highlighting glia as an active component of HD-like disease progression.

## Introduction

Huntington’s disease (HD) is an autosomal dominant neurodegenerative disorder caused by expansion of a cytosine–adenine–guanine (CAG) repeat in exon 1 of the ubiquitously expressed *huntingtin* (*HTT*) gene. The resulting polyglutamine-expanded mutant huntingtin (mHTT) progressively misfolds, oligomerises, and aggregates, disrupting multiple cellular pathways. HD manifests as progressive motor, cognitive, and psychiatric dysfunction, ultimately causing severe disability and premature death. Despite ubiquitous mHTT expression, degeneration preferentially affects striatal medium spiny neurons and specific cortical populations (Bates, 2025; Donaldson et al., 2026), suggesting that disease progression depends not only on mHTT but also on the cellular context in which it is processed.

HD research has traditionally focused on the well-established cell-autonomous toxicity of neuronal mHTT, which disrupts proteostasis, mitochondrial function, axonal transport, transcription, and synaptic integrity (Bhat et al., 2024; Ojalvo-Pacheco et al., 2024; Tong et al., 2024). However, neuronal survival depends on glia, which regulate ion balance, neurotransmitter recycling, metabolic and trophic support, synaptic integrity, and immune surveillance (Rangel-Gomez et al., 2024; Wen & Turrigiano, 2024). Because HTT is expressed in both neurons and glia, glia-originating pathology may contribute beyond established neuron-autonomous effects. HD may therefore involve dysfunction of the neuron–glia unit, with glial pathology and altered intercellular interactions contributing to progressive neurodegeneration (Jansen et al., 2017; Villanueva et al., 2025; Wilton & Stevens, 2020).

Glia also maintain proteostasis by recognizing, internalizing, and degrading extracellular protein aggregates through endosomal, lysosomal, and autophagic pathways (Gao et al., 2023; Wilson et al., 2023), buffering neurons against proteotoxic stress in HD and related polyglutamine disorders (Cvetanovic & Gray, 2023; Li & Haney, 2020). Persistent accumulation of aggregation-prone proteins, however, can overwhelm these processes, impair clearance, promote intracellular aggregation, and compromise glial support of neuronal homeostasis, potentially converting glia from protective scavengers into contributors to disease progression (Gao et al., 2023; Wilton & Stevens, 2020).

Consistent with this dual role, glial expression of polyglutamine-expanded proteins, α-synuclein, tau, and amyloid-β can induce behavioural deficits and non-cell-autonomous neuronal dysfunction in *Drosophila* and other models (Cvetanovic & Gray, 2023; Wilton & Stevens, 2020). mHTT toxicity may also extend beyond its expressing cells through mechanisms including extracellular vesicles, tunnelling nanotubes, synaptic release, and glial phagocytic pathways (Babcock & Ganetzky, 2015; Donnelly et al., 2020; Tarasiuk & Scuteri, 2022; Victoria & Zurzolo, 2017). These findings challenge a strictly cell-autonomous model and raise the possibility that glia-initiated pathology may progressively involve neuronal compartments (Chang et al., 2024; Goldman, 2024; Wilton & Stevens, 2020).

A critical unresolved question is whether mHTT pathology originating in glia can itself establish a progressive pathological sequence culminating in neuronal involvement, or whether glial alterations primarily represent compensatory or secondary responses to neuron-driven disease. Although neuronal mHTT toxicity is well established, glia are increasingly implicated in mHTT handling, intercellular aggregate dynamics, and neuronal homeostasis. Defining the temporal sequence initiated by glial mHTT is therefore essential for determining its contribution to HD progression and potential therapeutic relevance.

We addressed this question in *Drosophila melanogaster* using pan-glial expression of human polyglutamine-expanded Huntingtin exon 1. Longitudinal behavioural analyses were integrated with quantitative immunohistochemistry of mHTT accumulation, fluorescence intensity, aggregate architecture, and cell-type-specific localization to define the progression of glial pathology and accompanying neuronal involvement. Pan-neuronal expression of the identical mHTT construct using *elav*-GAL4 provided an established neuron-initiated reference for interpreting the temporal features of glia-initiated pathology.

We hypothesized that glia-originating mHTT can establish a progressive disease trajectory and ultimately influence neuronal function. Using neuronal mHTT expression as an established reference, we examined proteotoxic burden, aggregate dynamics, cellular association, and functional decline over time. This framework enables assessment of how glial pathology and neuron–glia interactions contribute to HD progression and may identify glia as a therapeutic target.

## Results

### Glial mHTT expression induces early functional abnormalities followed by progressive behavioural decline

Glial mHTT expression produced developmental and behavioural dysfunction, with the established neuronal phenotype providing a reference for glia-initiated pathology (Fig. 1). Third-instar *Repo*>HTTex1pQ93 larvae crawled approximately 1.6 cm versus 2.6 cm in *Repo*>HTTex1pQ25 controls, an approximately 38% reduction, whereas *elav*>HTTex1pQ93 larvae crawled approximately 0.45 cm versus 2.8 cm in *elav*>HTTex1pQ25 controls, an approximately 84% reduction (Fig. 1A). Thus, glial mHTT produced detectable, but comparatively moderate, larval locomotor dysfunction.

**Figure 1.**
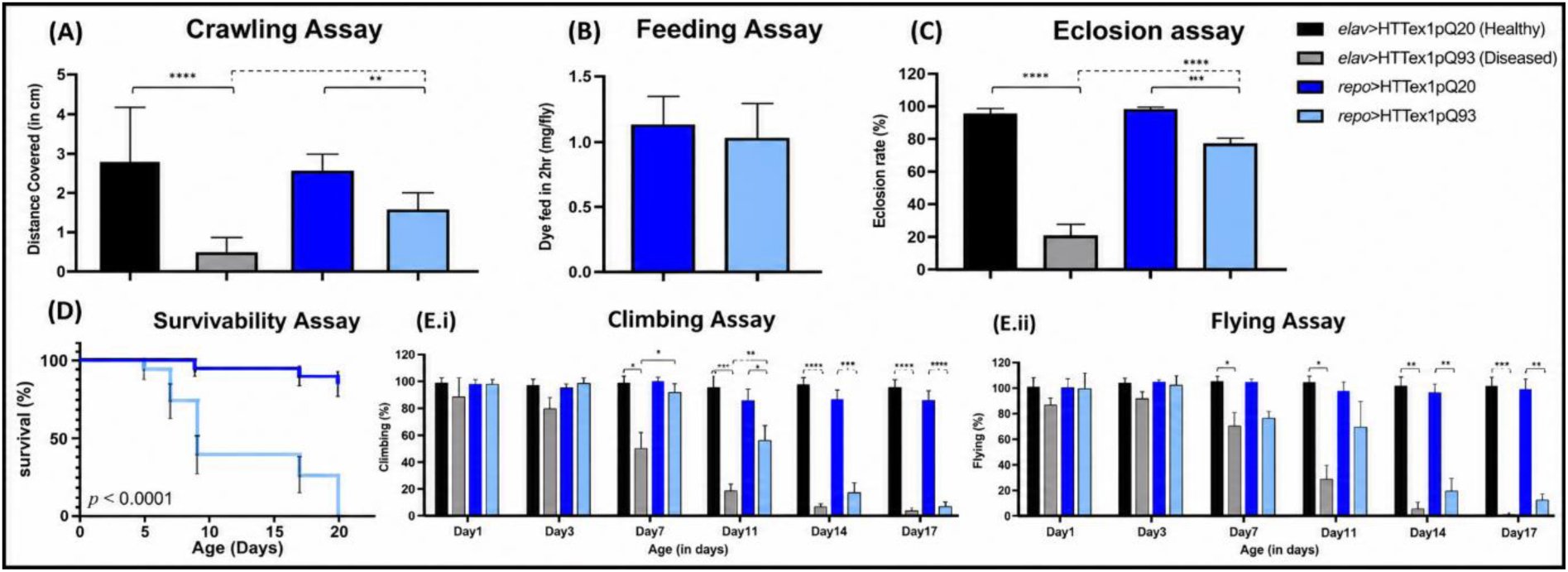
Effect of glial (*repo-*Gal4) and neuronal (*elav*-Gal4) expression of HTTex1pQ93 (mHTT) on (A) Crawling, (B) Feeding, **(C) Eclosion rate, (D) Survivability, and (E) Adult Locomotory functions.** (A) Glial expression of mHTT significantly decreased the crawling ability compared to age matched control (*repo*>HTTex1pQ20). The mobility decline was however less significant, compared to neuronal expression of mHTT; (B) Glial expression of mHTT do not compromise larval feeding behavior, compared to control (*repo*>HTTex1pQ20); (C) Glial expression of mHTT decreases eclosion rate less significantly compared to neuronal expression of mHTT; (D) Median survival of flies expressing mHTT in glial cells is compromised and 50% death is noted by 9 days post eclosion whereas upon neuronal expression of mHTT, 50% death is reported within 5 days post eclosion (Kumari et al., 2022); (E) Glial expression of mHTT significantly decreases (E.i.) climbing and (E.ii.) flying ability of flies compared to age matched control (*repo*>HTTex1pQ20). The decline in motor functions however has a delayed onset, and is less significant, compared to age matched flies having neuronal expression of mHTT. Data are presented as mean ± SEM unless otherwise indicated. Statistical analyses were performed as follows: (A, C) Kruskal– Wallis test followed by Tukey’s multiple-comparison test; (B) unpaired Student’s t-test; (D) Kaplan–Meier survival analysis with log-rank (Mantel–Cox) test; (E) two-way ANOVA followed by Tukey’s multiple-comparison test. For panel (E), significance symbols denote the following comparisons: *, *elav*>HTTex1pQ93 versus *repo*>HTTex1pQ93; #, *repo*>HTTex1pQ25 versus *repo*>HTTex1pQ93; +, *elav*>HTTex1pQ25 versus *elav*>HTTex1pQ93. Statistical significance is indicated as P < 0.05, P < 0.01, P < 0.001, and P < 0.0001. Sample sizes were: (A) n = 40 larvae per condition (10 larvae/replicate, 3 technical trials per replicate); (B) n = 40 larvae per condition (3 biological replicates); (C) 100 eggs per replicate (5 biological replicates); (D) n = 100 flies per genotype (20 flies/replicate, 5 biological replicates); (E) n = 30 flies per genotype and time point (10 flies/replicate, 3 biological replicates).

This deficit was not accompanied by altered feeding. *Repo*>HTTex1pQ93 larvae consumed approximately 1.0 mg dye h⁻¹ versus approximately 1.1 mg h⁻¹ in *Repo*>HTTex1pQ25 controls, with no significant difference between genotypes (Fig. 1B). Thus, reduced locomotion occurred without detectable reduction in food intake.

Developmental effects were also evident during eclosion (Fig. 1C). Approximately 77% of *Repo*>HTTex1pQ93 animals eclosed versus 97% of *Repo*>HTTex1pQ25 controls, an approximately 21% reduction, with developmental arrest at different pupal stages. In the neuronal reference, approximately 20% of *elav*>HTTex1pQ93 animals eclosed versus 95% of *elav*>HTTex1pQ25 controls, an approximately 79% reduction. Together with the crawling phenotype, these findings establish developmental and functional abnormalities following glial mHTT expression before adulthood, although less severe than with direct neuronal expression.

Glial mHTT subsequently produced progressive adult mortality (Fig. 1D). *Repo*>HTTex1pQ25 flies maintained approximately 90% survival through Day 20, whereas *Repo*>HTTex1pQ93 survival declined to approximately 75% by Day 7, 40% by Day 9, and 25% by Day 17, with complete mortality by approximately Day 20 (P < 0.0001) and median survival of approximately 9 days. Previous neuronal mHTT analysis showed approximately 50% mortality by Day 5. Thus, glial mHTT produced developmental abnormalities followed by delayed but progressive adult mortality relative to the rapid neuronal phenotype.

This progression was particularly evident in adult locomotor behaviour. *repo*>HTTex1pQ93 flies retained approximately 96–98% climbing ability through Day 3 and 90% at Day 7, declining to approximately 56% at Day 11 and 13% at Day 14, with near-complete loss by Day 17 (Fig. 1E.i); *repo*>HTTex1pQ25 controls maintained approximately 86–99%. In the neuronal reference, *elav*>HTTex1pQ93 climbing declined from approximately 87% at Day 1 and 79% at Day 3 to 50% at Day 7, 16% at Day 11, and 10% at Day 14, with essentially no climbing by Day 17; *elav*>HTTex1pQ25 controls maintained approximately 93–99%. Thus, glial mHTT preserved early adult climbing before pronounced deterioration, with the major decline occurring approximately 4 days later than in the neuronal condition.

Flight performance followed the same pattern (Fig. 1E.ii). *repo*>HTTex1pQ93 flies retained approximately 93–97% flight ability during Days 1–3, declining to 77% at Day 7, 67% at Day 11, 20% at Day 14, and 4% at Day 17; *repo*>HTTex1pQ25 controls maintained approximately 93–100% across most of the analysed period. *elav*>HTTex1pQ93 flies retained approximately 83–87% during Days 1–3, declining to approximately 70% at Day 7, 30% at Day 11, and 7% at Day 14, with flight essentially lost shortly before terminal mortality by Day 15; *elav*>HTTex1pQ25 controls similarly maintained approximately 93–100%. Thus, preserved early flight following glial mHTT expression preceded substantial late deterioration.

Collectively, glial mHTT established a progressive phenotype comprising moderate developmental abnormalities, preserved early adult motor function, and subsequent deterioration in climbing, flight, and survival. Despite delayed progression, *repo*>HTTex1pQ93 flies ultimately showed near-complete loss of climbing and flight by Day 17. These findings define a temporal window in which glia-initiated pathology precedes pronounced functional deterioration, providing a basis for examining the accompanying cellular and aggregate-associated changes.

### Glial mHTT expression establishes progressive cellular pathology characterised by increasing HTT burden and altered glial immunoreactivity

Confocal imaging and quantitative fluorescence analysis revealed progressive cellular pathology following glial HTTex1pQ93 expression, with neuronal expression providing an established reference (Figs. 2–4). In both paradigms, HTTex1pQ20 controls showed minimal HTT immunoreactivity and broadly preserved neuronal and glial staining throughout the analysed period (Figs. 2A and 3A). In contrast, HTTex1pQ93 produced pronounced HTT accumulation with altered cell-type-specific marker immunoreactivity (Figs. 2B, 3B and 4). Representative images defined the spatial and morphological changes, while fluorescence measurements established their temporal profiles.

**Figure 2.**
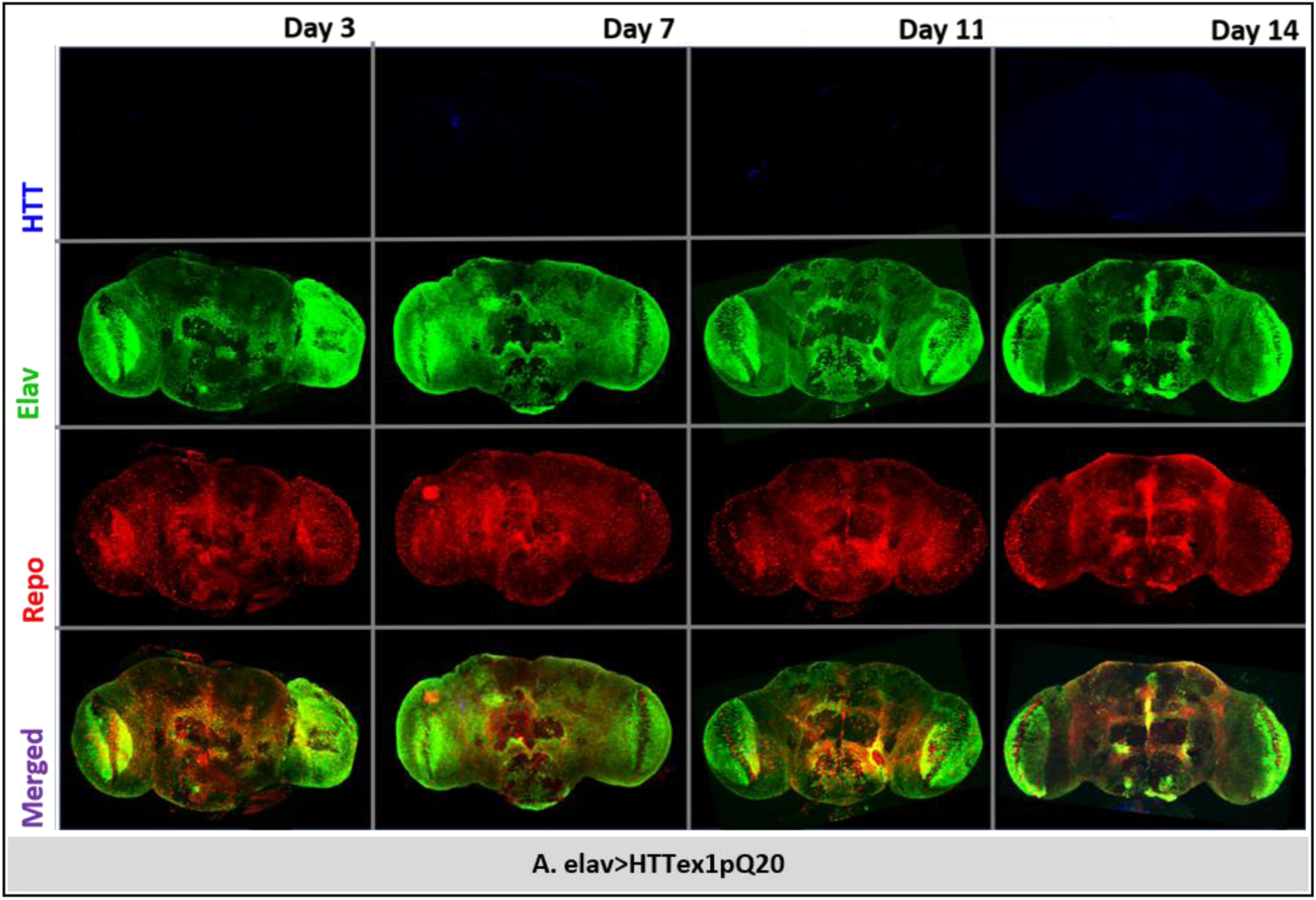
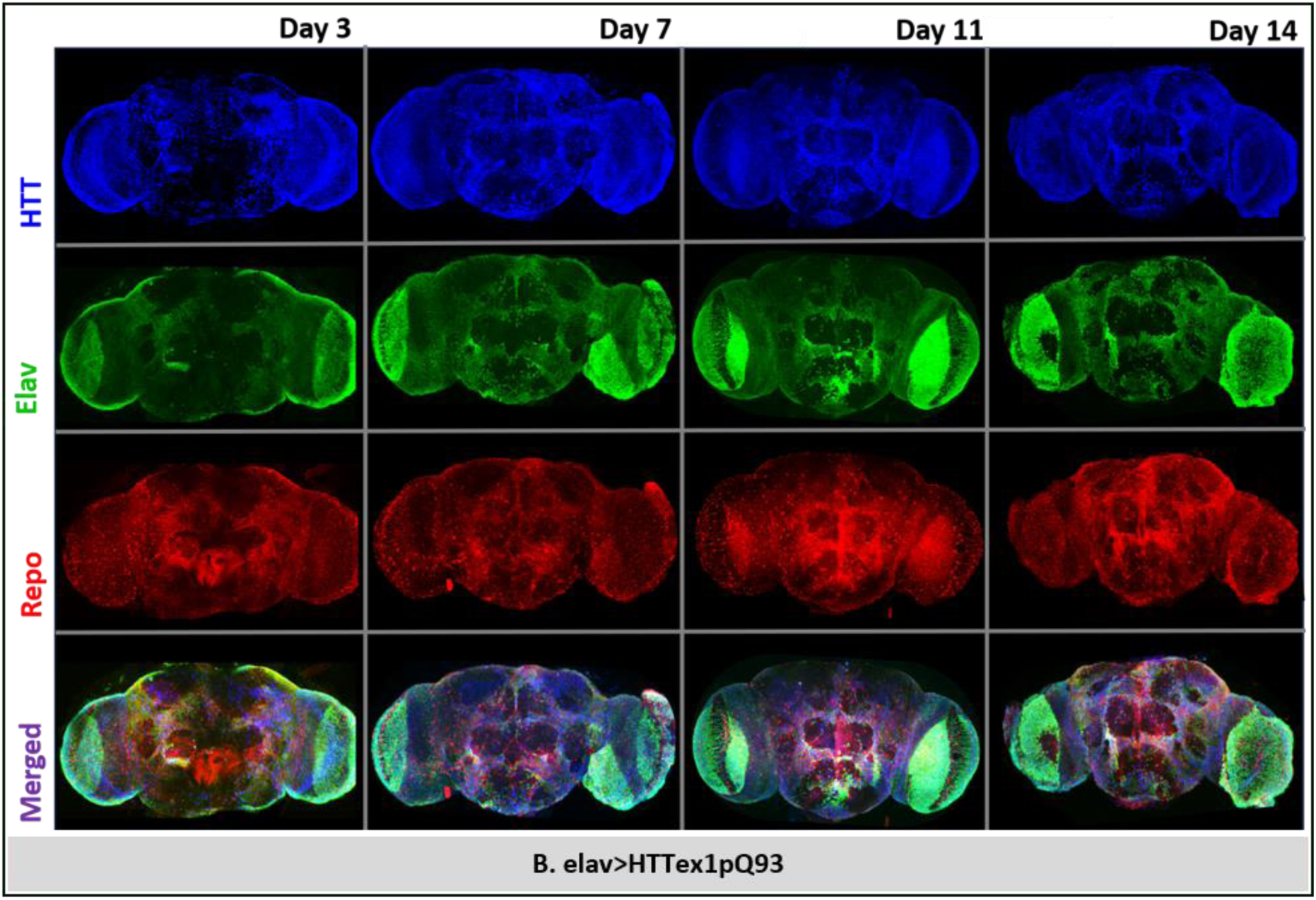
Early and progressive neuronal mutant huntingtin accumulation in *elav*>HttQ93 adult Drosophila brains. (A) Representative confocal maximum-intensity projection images of *elav*>HTTex1pQ20 (control) adult brains collected at Days 3, 7, 11, and 14 following eclosion. Brains were immunostained for mutant huntingtin (HTT, blue), *elav* (neuronal marker, green), and *repo* (glial marker, red). Control brains exhibited minimal HTT immunoreactivity together with preserved neuronal and glial staining patterns throughout the analysed period. (B) Representative confocal maximum-intensity projection images of *elav*>HTTex1pQ 93 adult brains at the corresponding ages.

**Figure 3.**
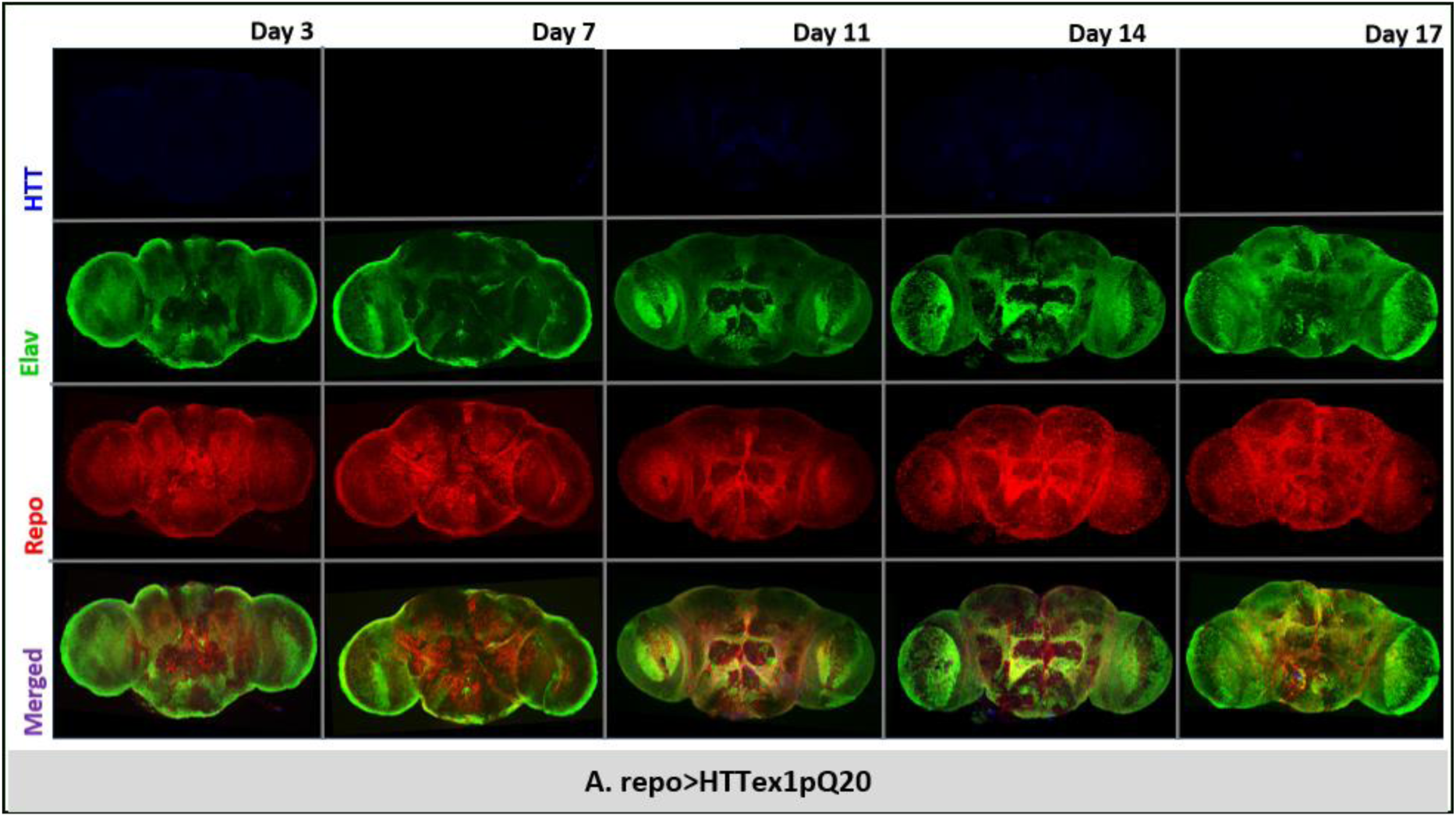
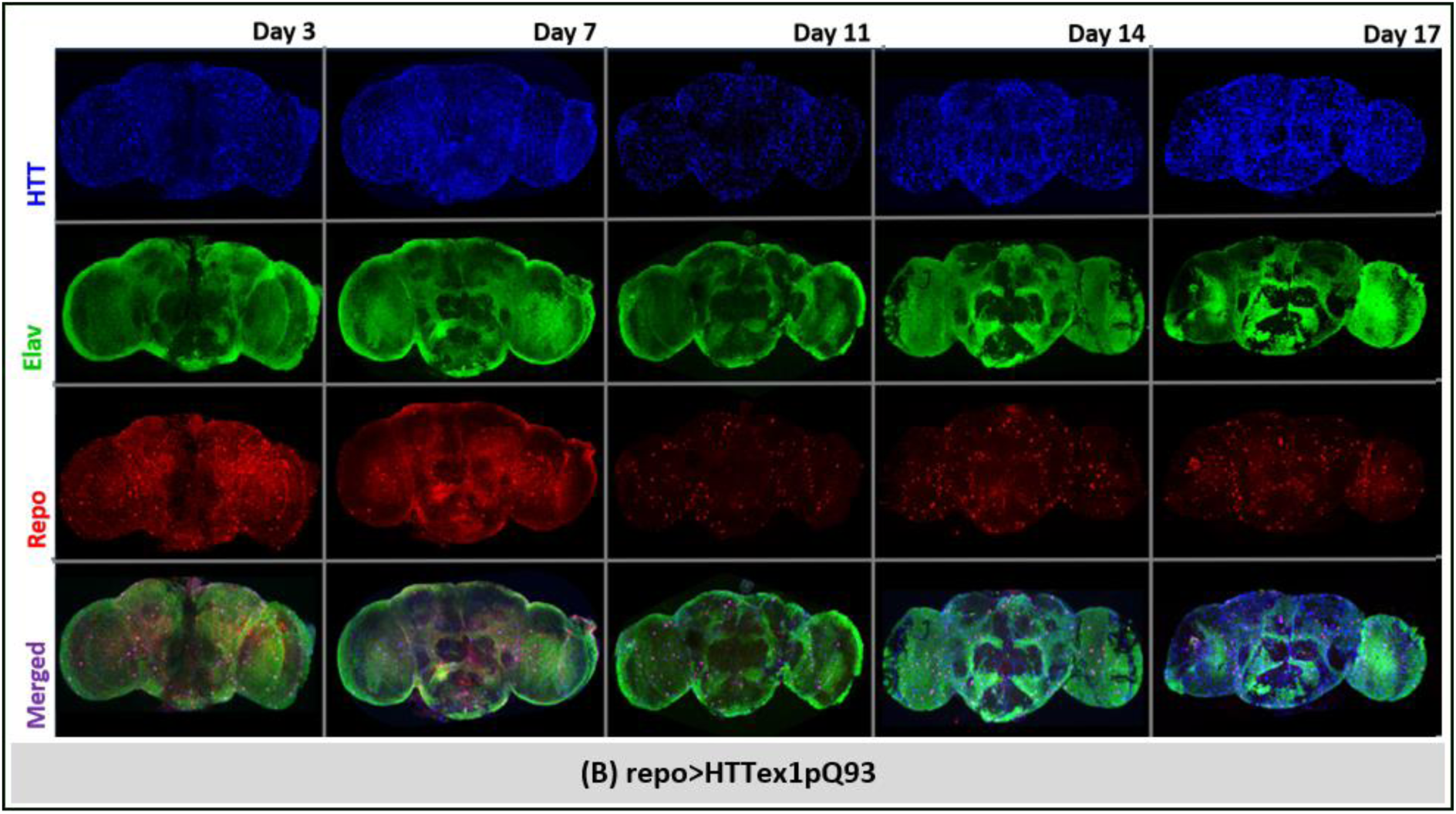
Progressive glial pathology and mutant huntingtin accumulation in *repo*>HttQ93 adult Drosophila brains. (A) Representative confocal maximum-intensity projections of *repo*>HTTex1pQ20 (control) adult brains collected at Days 3, 7, 11, 14, and 17 following eclosion. Brains were immunostained for mutant huntingtin (HTT, blue), *elav* (neuronal marker, green), and *repo* (glial marker, red). Control brains exhibited negligible HTT immunoreactivity together with preserved neuronal and glial staining patterns throughout the analysed period. (B) Representative confocal maximum-intensity projections of *repo*>HTTex1pQ93 adult brains at the corresponding ages.

**Figure 4.**
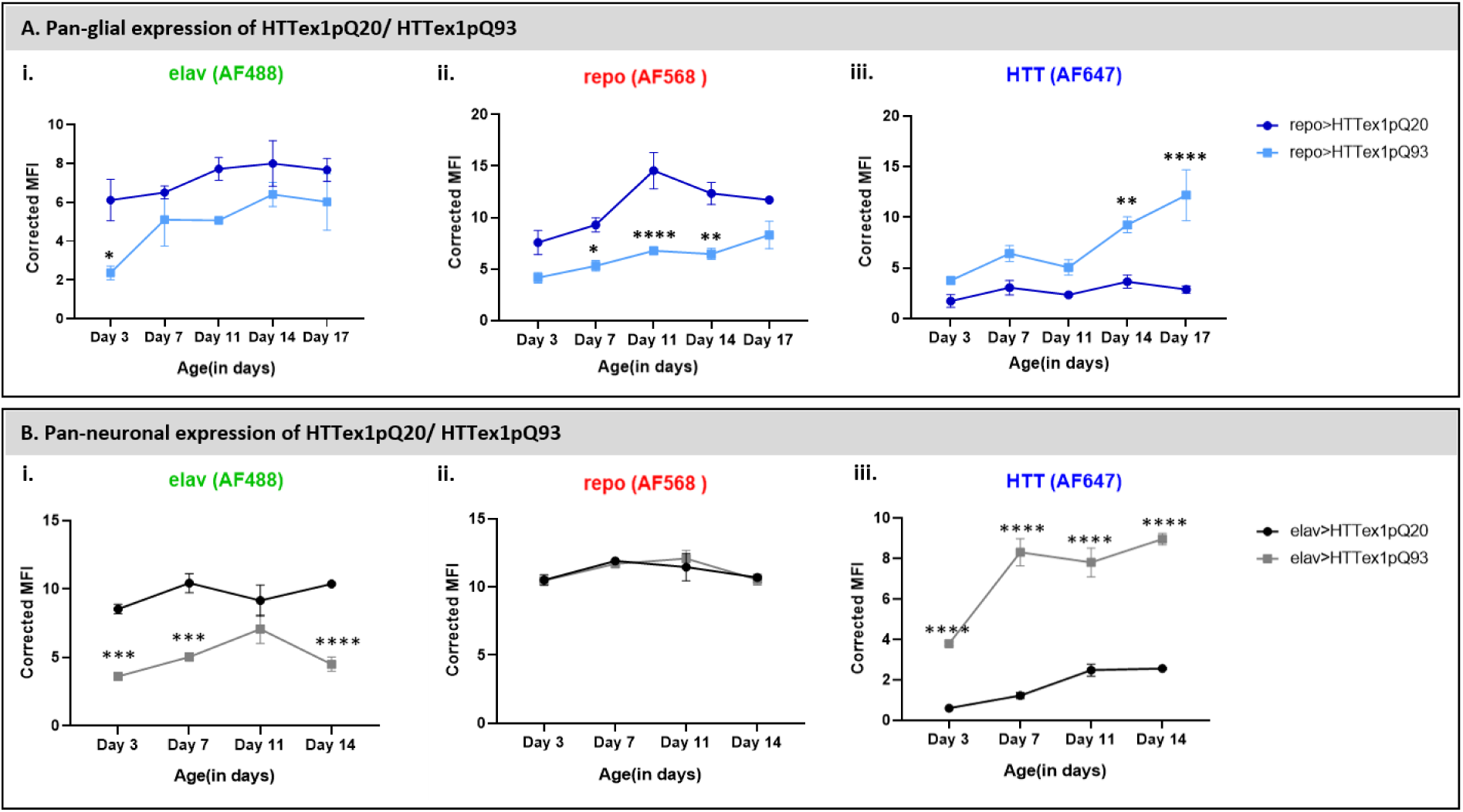
Temporal changes in neuronal, glial and HTT markers following neuronal or glial expression of mutant huntingtin. Corrected mean fluorescence intensity (MFI) of *elav* (Alexa-fluor 488), *repo* (Alexa-fluor 568) and mutant huntingtin (HTT; Alexa-fluor 647) immunoreactivity in adult brains having pan-glial (A) or pan neuronal (B) expression of HTTex1pQ20 and HTTex1pQ93; brains collected at Days 3, 7, 11, 14 and 17 post-eclosion. Corrected MFI was calculated by subtracting the background fluorescence intensity from the mean fluorescence intensity of each optical section and averaging the values across all Z-sections of each brain. Data are presented as mean ± SEM. Statistical significance was determined using two-way ANOVA followed by Šídák’s multiple-comparison test, comparing immunoreactivity of *elav*/*repo*/HTT markers in pan-glial or pan-neuronal expression of HTTex1pQ20 and HTTex1pQ93 at each age. *P < 0.05, **P < 0.01, ***P < 0.001 and ****P < 0.0001.

The neuronal reference showed early, persistent pathology. *elav*>HTTex1pQ20 brains exhibited minimal HTT immunoreactivity from Days 3–14, without conspicuous HTT-positive puncta, while *elav* and *repo* staining remained broadly distributed and comparatively stable (Fig. 2A). In contrast, *elav*>HTTex1pQ93 brains displayed abundant, widely distributed HTT immunoreactivity and numerous punctate structures from Day 3 through Days 7, 11, and 14 (Fig. 2B). Despite this burden, the gross anatomical distributions of both *elav* and *repo* immunoreactivity remained broadly preserved across the central brain and optic lobes.

Quantification revealed changes not readily apparent from representative images (Fig. 4B). Corrected *elav* mean fluorescence intensity (MFI) was significantly reduced in *elav*>HTTex1pQ93 brains relative to *elav*>HTTex1pQ20 controls at Day 3 (P < 0.0001), Day 7 (P < 0.001), and Day 14 (P < 0.0001), with a similar but non-significant reduction at Day 11 (Fig. 4B.i). In contrast, *repo* MFI remained comparable between genotypes throughout the analysed period (Fig. 4B.ii). Corrected HTT MFI was significantly elevated in *elav*>HTTex1pQ93 brains from Day 3 onward, with differences maintained at Days 3, 7, 11, and 14 (all P < 0.0001; Fig. 4B.iii). Thus, the neuronal reference was characterized by rapid, sustained HTT accumulation and reduced neuronal marker immunoreactivity, while glial *repo* immunoreactivity remained comparatively preserved (Figs. 2 and 4B).

In contrast, pan-glial mHTT expression produced progressively evolving cellular pathology. *repo*>HTTex1pQ20 brains showed minimal HTT immunoreactivity from Days 3–17, with broadly preserved *repo* and *elav* staining across the central brain and optic lobes and no conspicuous HTT-positive puncta or major age-dependent changes (Fig. 3A). In *repo*>HTTex1pQ93 brains, HTT was detectable throughout the brain by Day 3, followed by increasingly prominent HTT-positive puncta and aggregates across the central brain and optic lobes with age (Fig. 3B). Concurrently, *repo* immunoreactivity became less diffuse and increasingly heterogeneous, with intensely labelled punctate structures against a less uniform background, particularly at Days 11–17. Merged images showed increasing spatial association of HTT-positive aggregates with neuronal structures. Nevertheless, *elav*-positive neuronal structures remained readily identifiable, and overall brain morphology was preserved even at later stages.

Quantitative fluorescence analysis confirmed progressive glial alteration (Fig. 4A). *repo* MFI was lower in *repo*>HTTex1pQ93 than *repo*>HTTex1pQ20 brains throughout much of the time course, reaching significance at Day 7 (P < 0.05), Day 11 (P < 0.0001), and Day 14 (P < 0.01; Fig. 4A.ii), paralleling the increasingly heterogeneous *repo* staining. In contrast, *elav* MFI was significantly reduced only at Day 3 (*P* < 0.05), with no significant differences thereafter (Fig. 4A.i), consistent with comparatively preserved neuronal staining during progressive glial pathology.

HTT immunoreactivity increased during glial disease progression (Fig. 4A.iii). *repo*>HTTex1pQ93 brains showed higher HTT MFI than *repo*>HTTex1pQ20 controls throughout the analysed period, reaching significance at Day 11 (P < 0.01) and Day 17 (P < 0.0001). Thus, increasing HTT burden accompanied progressive alteration and reduction of *repo* immunoreactivity.

Despite these cellular changes, gross brain morphology remained largely preserved, as in the neuronal reference (Figs. 2 and 3). The central brain and optic lobes remained identifiable without obvious large-scale tissue collapse, indicating that progression predominantly involved protein accumulation and altered cellular immunoreactivity rather than overt disruption of gross brain architecture within the examined period.

Together, glial mHTT expression produced a progressive cellular sequence characterized by increasing HTT burden and declining, increasingly heterogeneous *repo* immunoreactivity, while overall *elav*-positive neuronal staining remained comparatively preserved (Figs. 3 and 4A). In the neuronal reference, HTT accumulation was already pronounced at the earliest analysed age and accompanied by reduced *elav* immunoreactivity, while *repo* signal remained largely unchanged (Figs. 2 and 4B). Thus, glial mHTT defines a period of evolving glial pathology with preserved gross neuronal architecture, providing the basis for subsequent analyses of aggregate architecture and progressive HTT association with neuronal and glial compartments.

### Glial mHTT pathology is accompanied by progressive neuronal HTT association

To determine how mHTT association evolves following glial expression, thresholded Manders’ (M2) coefficient quantified the fraction of HTT signal overlapping *elav*-positive neuronal or *repo*-positive glial compartments (Fig. 5). Neuronal mHTT expression provided an established reference for HTT distribution when originating within neurons.

**Figure 5.**
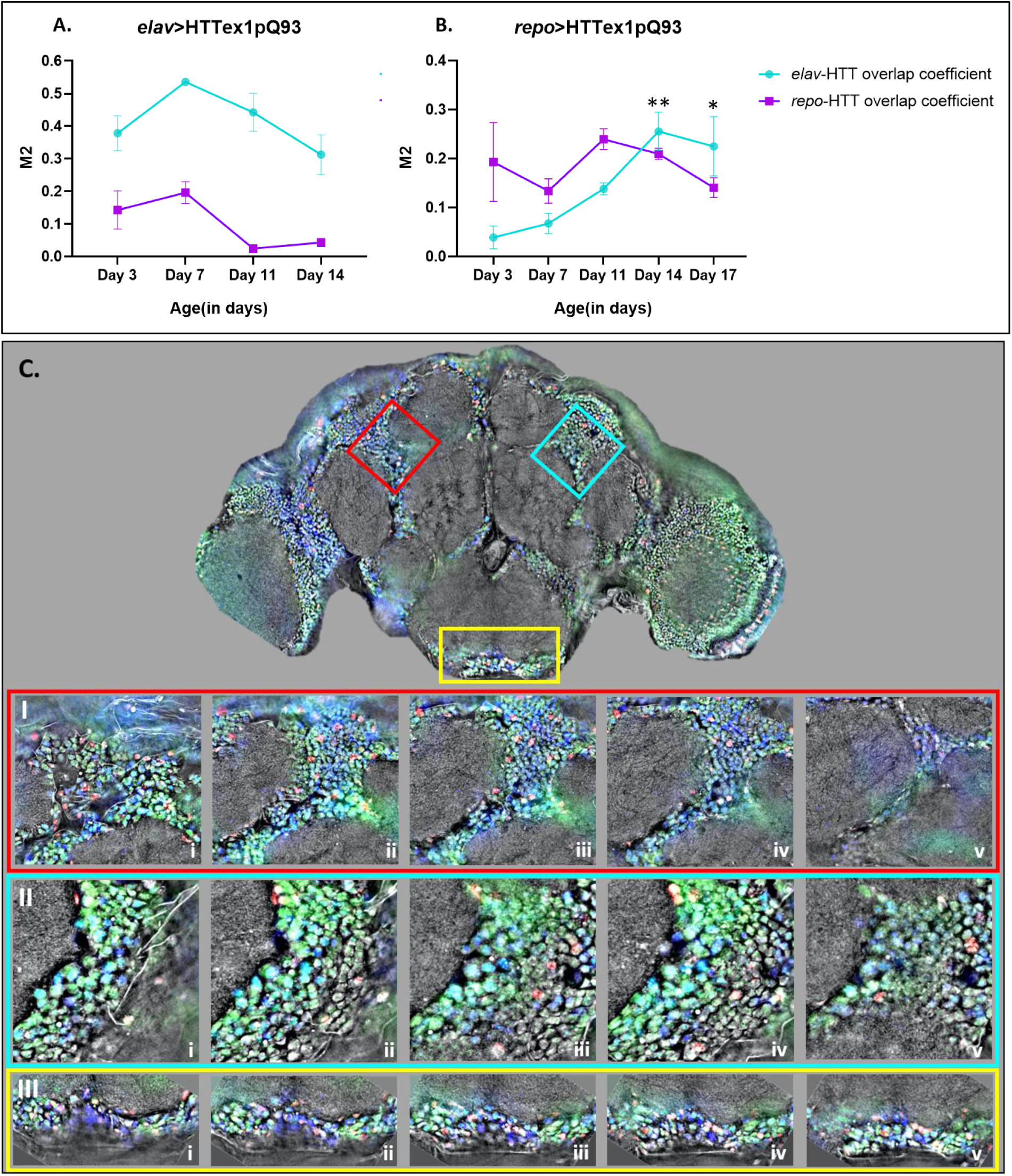
Age-dependent changes in neuronal and glial association of mutant huntingtin. (A–B) Thresholded Manders’ M2 coefficient quantifying the fraction of HTT signal overlapping *elav*-positive neuronal (cyan) or *repo*-positive glial (magenta) compartments in *elav*>HTTex1pQ93 (A) and *repo*>HTTex1pQ93 (B) adult brains. In *elav*>HTTex1pQ93 brains, two-way ANOVA revealed significant effects of cell type (F(1,16) = 107.3, P < 0.0001) and age (F(3,16) = 6.718, P = 0.0038), but no age × cell-type interaction (F(3,16) = 1.754, P = 0.1965), indicating persistent preferential neuronal HTT association without a significant age-dependent shift between compartments. Tukey’s multiple-comparison test, using Day 3 as reference within each colocalization coefficient, detected no significant differences at subsequent time points. In *repo*>HTTex1pQ93 brains, age (F(4,20) = 4.175, P = 0.0128) and age × cell-type interaction (F(4,20) = 3.495, P = 0.0255) were significant, whereas the overall cell-type effect was not (F(1,20) = 2.541, P = 0.1266), indicating an age-dependent change in relative neuronal and glial HTT association. *elav*–HTT M2 increased significantly at Day 11 (P < 0.01) and Day 17 (P < 0.05) relative to Day 3, whereas *repo*– HTT M2 showed no significant age-dependent change. (C) Representative correlative holotomography and fluorescence imaging of an intact Day 14 *repo*>HTTex1pQ93 adult brain. The Holo-CT image provides label-free structural context overlaid with *elav* (green), *repo* (red), and HTT (blue) fluorescence. Three regions of interest (red, cyan, and yellow boxes) in the optimal whole-brain Z-plane are shown as sequential Z-plane montages I, II, and III (i–v), respectively. HTT-positive structures show close spatial association with *elav*-positive neuronal compartments across successive Z-planes, with cyan/blue-green overlap indicating neuronal HTT association despite glial-directed mHTT expression. This representative three-dimensional observation provides qualitative spatial support for the increased *elav*–HTT association detected by longitudinal M2 analysis but does not establish direct glia-to-neuron mHTT transfer. Data are mean ± SEM. Two-way ANOVA assessed effects of age and cell type, followed by Tukey’s multiple-comparison test comparing each time point with Day 3 within the respective colocalization coefficient. *P < 0.05, **P < 0.01.

In *elav*>HTTex1pQ93 brains, HTT consistently showed greater overlap with *elav*-positive neuronal than *repo*-positive glial compartments throughout Days 3–14 (Fig. 5A). Two-way ANOVA revealed significant effects of cell type (F(1,16) = 107.3, P < 0.0001) and age (F(3,16) = 6.718, P = 0.0038), but no age × cell-type interaction (F(3,16) = 1.754, P = 0.1965), indicating that relative neuronal versus glial association did not significantly change with age. Consistently, Tukey’s multiple-comparison analysis using Day 3 as reference detected no significant age-dependent changes in either *elav*–HTT or *repo*–HTT M2. Thus, neuronally expressed mHTT remained preferentially associated with neuronal compartments throughout the analysed period.

In contrast, glial mHTT showed progressively changing cellular association (Fig. 5B). At Day 3, HTT overlap was greater with *repo*-positive glial than *elav*-positive neuronal compartments, consistent with glial-targeted expression. With age, *elav*–HTT M2 increased, whereas *repo*–HTT M2 showed no significant change relative to Day 3. Two-way ANOVA revealed a significant effect of age (F(4,20) = 4.175, P = 0.0128) and age × cell-type interaction (F(4,20) = 3.495, P = 0.0255), while the overall cell-type effect was not significant (F(1,20) = 2.541, P = 0.1266). Thus, the relative neuronal and glial association of HTT changed during glia-initiated pathology.

Post hoc analysis localized this change to the neuronal compartment. Relative to Day 3, *elav*–HTT M2 increased significantly at Day 11 (*P* < 0.01) and Day 17 (*P* < 0.05), whereas *repo*–HTT M2 remained statistically unchanged across ages (Fig. 5B). Thus, mHTT was initially preferentially associated with glial compartments, but disease progression was accompanied by increasing neuronal HTT association without a corresponding age-dependent increase in *repo*–HTT overlap.

Correlative Holo-CT and fluorescence imaging of a representative *repo*>HTTex1pQ93 brain at Day 14 further examined this spatial relationship (Fig. 5C). HTT-positive structures were closely associated with *elav*-positive neuronal compartments, with some appearing localised within *elav*-positive domains across successive Z-planes. This provides qualitative three-dimensional anatomical support for the increased *elav*–HTT association detected by longitudinal Manders’ M2 analysis. As Holo-CT imaging was performed at a single time point, it supports spatial association without establishing temporal redistribution or direct glia-to-neuron mHTT transfer.

Collectively, these analyses identify increasing neuronal HTT association as a progressive feature of glia-initiated mHTT pathology. Glial mHTT was initially preferentially associated with *repo*-positive compartments, followed by increasing *elav*–HTT association with age despite unchanged *repo*–HTT overlap (Fig. 5). In contrast, the neuronal reference remained preferentially neuron-associated without significant age-dependent redistribution between compartments. Holo-CT imaging provides complementary spatial support for the quantitative M2 findings, while neither analysis establishes temporal redistribution or direct glia-to-neuron transfer of mHTT.

### Neuronal and glial mHTT expression exhibit distinct temporal patterns of aggregate remodelling

Quantitative analysis of aggregate number and volume defined how mHTT populations evolved following glial and neuronal expression (Fig. 6). Aggregates were classified by volume as puncta, small aggregates, large aggregates, or inclusions, enabling longitudinal analysis of aggregate composition.

**Figure 6.**
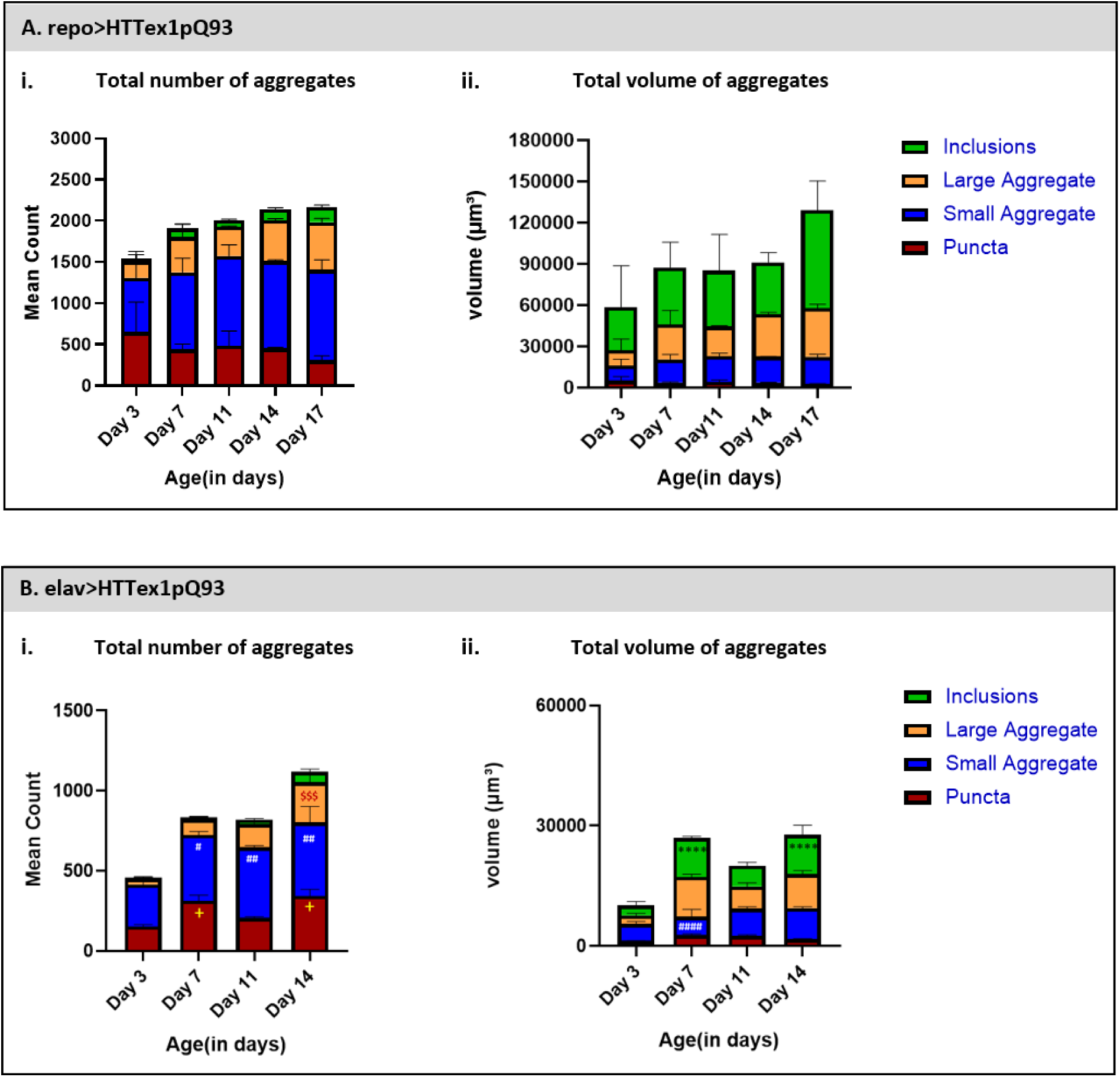
Quantitative analysis of aggregate number and volume composition following glial and neuronal expression of mutant huntingtin. (A) Aggregate composition in *repo*>HTTex1pQ93 adult brains during disease progression. The left panel shows the total number of aggregates per brain, while the right panel shows the total aggregate volume (μm³). Stacked bars represent the contribution of puncta (red), small aggregates (blue), large aggregates (orange), and inclusions (green) at Days 3, 7, 11, 14 and 17 post-eclosion. (B) Aggregate composition in *elav*>HTTex1pQ93 adult brains. The left panel shows the total number of aggregates per brain, while the right panel shows the total aggregate volume (μm³). Aggregate classes are represented using the same colour scheme as in panel A and were analysed at Days 3, 7, 11 and 14 post-eclosion. Aggregate classes were defined using fixed percentile-derived volume thresholds generated from the pooled global distribution of aggregate volumes and were applied uniformly to all brains irrespective of genotype or age. Data are presented as mean ± SEM (n = 3 biologically independent brains per time point). For both aggregate count and aggregate volume, statistical analyses were performed using ordinary two-way ANOVA with age and aggregate class as independent variables. Post hoc comparisons were performed using Tukey’s multiple-comparison test, with Day 3 serving as the reference for each aggregate class independently. Statistical symbols indicate comparisons within the corresponding aggregate class only: +, puncta; #, small aggregates; $, large aggregates; and *, inclusions. The number of symbols denotes the level of statistical significance (*P* < 0.05, P < 0.01, *P* < 0.001 and P < 0.0001). Aggregate number and volume analyses demonstrate distinct temporal aggregation profiles in the two models, with *repo*>HTTex1pQ93 exhibiting relatively stable aggregate composition during progressive accumulation, whereas *elav*>HTTex1pQ93 displays more pronounced temporal remodelling of aggregate populations.

In *repo*>HTTex1pQ93 brains, total aggregate number increased from Days 3–7 and remained comparatively stable through Day 17, with modest changes among aggregate classes (Fig. 6A.i). Aggregate volume varied over time and was greatest at Day 17 (Fig. 6A.ii). Two-way ANOVA identified a significant effect of aggregate class (F(3,40) = 11.60, P < 0.0001), but not age (F(4,40) = 1.89, P = 0.1309) or age × aggregate class interaction (F(12,40) = 0.88, P = 0.5775). Thus, after early establishment, the relative volume distribution of glial aggregate populations remained comparatively stable without pronounced age-dependent remodelling.

In contrast, neuronal mHTT produced marked temporal remodelling (Fig. 6B). Total aggregate number increased substantially from Days 3–7 and continued changing through Days 11 and 14, with aggregate classes showing distinct temporal profiles (Fig. 6B.i). Aggregate volume increased markedly by Day 7, decreased at Day 11, and increased again by Day 14 (Fig. 6B.ii). Two-way ANOVA revealed significant effects of aggregate class (F(3,32) = 21.05, P < 0.0001), age (F(3,32) = 18.83, P < 0.0001), and age × aggregate class interaction (F(9,32) = 4.32, P = 0.0010), demonstrating differential temporal changes among aggregate classes. Thus, neuronal mHTT was characterized by continuous restructuring of aggregate architecture.

These analyses therefore revealed distinct aggregate trajectories. Following glial expression, aggregate number increased from Days 3–7 but remained comparatively stable thereafter, while the absence of significant age or age × aggregate class effects indicated no pronounced age-dependent remodelling of relative volume distribution among puncta, small aggregates, large aggregates, and inclusions. Conversely, neuronal mHTT showed significant age-dependent and class-specific changes in aggregate volume. Thus, glial pathology progressed within a comparatively stable aggregate population after early establishment, whereas neuronal pathology involved continued aggregate remodelling.

Integration with MFI analysis further defined mHTT accumulation within these populations. In *repo*>HTTex1pQ93 brains, HTT immunoreactivity progressively increased despite comparatively stable aggregate numbers after early establishment and no significant age-dependent remodelling of aggregate composition (Figs. 4A.iii and 6A). This pattern is consistent with continued mHTT accumulation through growth of established aggregate populations, potentially from pre-existing mHTT ‘seeds’, rather than sustained formation of new aggregates. In contrast, neuronal HTT accumulation accompanied marked age-dependent changes in aggregate number and volume composition (Figs. 4B.iii and 6B), with the significant age × aggregate class interaction demonstrating differential remodelling among aggregate populations. Together, these findings demonstrate that glial mHTT pathology progresses through increasing HTT burden within comparatively stable aggregate populations, whereas neuronal mHTT undergoes continued restructuring of the aggregate landscape, identifying cellular context as an important determinant of aggregate evolution during disease progression.

## Discussion

### Glial mHTT produces HD-like pathology with a distinct temporal trajectory from neuronal mHTT

The present study reveals that the pathogenic consequences of mHTT are strongly shaped by the cellular context in which it is expressed. Rather than producing simply a slower version of neuronal mHTT pathology, mHTT within glia appears to generate a distinct and ultimately severe pathological trajectory. Neuronal expression produced rapid intracellular mHTT accumulation, dynamic aggregate remodelling and early behavioural dysfunction. In contrast, glial expression initiated a slower sequence characterized by progressive mHTT accumulation, relatively stable aggregate architecture, declining *Repo* immunoreactivity, increasing neuronal HTT association and delayed behavioural deterioration. Thus, mHTT is not benign or merely less pathogenic in glia; its pathogenic consequences emerge through a different route from those of neuronal mHTT.

This distinction can be understood within the framework of cell-autonomous and non-cell-autonomous HD pathology. In neurons, mHTT directly disrupts proteostasis, transcriptional regulation, mitochondrial function, axonal transport and synaptic signalling, rapidly compromising neuronal and circuit function. In glia, mHTT initially perturbs the cellular systems responsible for maintaining the neural environment, including proteostasis, aggregate handling, metabolism and homeostatic functions. The resulting pathology may progressively alter neuron–glia interactions, allowing neuronal dysfunction to emerge despite mHTT originating outside neurons (Creus-Muncunill & Ehrlich, 2019; Donaldson et al., 2026; Goldman, 2024; Villanueva et al., 2025). Thus, the cellular site of mHTT expression determines the route through which the same pathogenic protein compromises the nervous system.

Evidence from other proteinopathies supports this context-dependent pathogenicity. Polyglutamine-expanded proteins expressed in *Drosophila* glia disrupt blood–brain barrier integrity through glia-specific proteostatic dysfunction (Yeh et al., 2018). Glial α-synuclein induces transcriptional programmes distinct from neuronal α-synuclein and promotes non-cell-autonomous neurodegeneration (Olsen & Feany, 2019), while glial tau primarily disrupts glia–axon interactions and axonal maintenance (Scarpelli et al., 2019). Together with evidence that glia differentially recognize, process and clear aggregation-prone proteins, these findings suggest that cell-type-specific differences in proteostatic capacity, trafficking, autophagy–lysosomal function, aggregate sequestration and metabolic homeostasis can shape both aggregate evolution and functional consequences (Li & Haney, 2020; Wilson et al., 2023; Chang et al., 2024; Cvetanovic & Gray, 2023).

Importantly, the delayed behavioural phenotype following glial mHTT expression should therefore not be equated with reduced pathogenicity. Instead, it may reflect a fundamentally different mode of toxicity. Neuronal mHTT produces relatively direct cell-autonomous dysfunction, whereas glial mHTT progressively compromises the cellular environment on which neuronal function depends. The initially slower phenotype may reflect compensatory glial homeostatic and proteostatic mechanisms that become increasingly insufficient as mHTT accumulates. This provides a framework in which glial mHTT can be catastrophic not because it immediately destroys glial function, but because its pathology progressively destabilizes the neuron–glia system, ultimately producing severe functional consequences (Creus-Muncunill & Ehrlich, 2019).

### Cell-type-specific proteostasis shapes mHTT aggregation and disease progression

A central finding of this study is that neuronal and glial mHTT expression produced distinct patterns of aggregate evolution. Neuronal expression caused rapid intracellular mHTT accumulation with significant age-dependent changes in aggregate number and volume, indicating continuous remodelling during disease progression. In contrast, glial expression produced progressive intracellular mHTT accumulation despite comparatively stable aggregate architecture after its initial establishment. Thus, neurons and glia appear to process the same pathogenic protein differently rather than merely accumulating it at different rates.

Proteostasis is increasingly recognized as a determinant of selective vulnerability in HD. Although mHTT is ubiquitously expressed, neurons and glia differ substantially in protein quality-control systems, including molecular chaperones, autophagy, lysosomal degradation, intracellular trafficking, and metabolic regulation ((Abjean et al., 2023; Benraiss et al., 2021; Jansen et al., 2017)). Single-cell transcriptomic studies further demonstrate cell-type-specific differences in stress-response and proteostatic networks during HD progression ((Al-Dalahmah et al., 2020)), supporting distinct modes of intracellular mHTT handling.

In neurons, the significant age × aggregate class interaction showed that aggregate populations remained dynamic, with different classes contributing unequally to total aggregate burden over time. Together with increasing HTT immunoreactivity and early behavioural decline, this suggests continuous nucleation, growth, remodelling, and clearance rather than a stable equilibrium. Glia showed the opposite pattern: aggregate architecture remained comparatively stable despite continued mHTT accumulation. Aggregate class remained the principal determinant of composition, whereas neither age nor the age × aggregate class interaction significantly influenced aggregate distribution. HTT immunoreactivity progressively increased while aggregate numbers changed relatively little after their initial establishment, suggesting that increasing mHTT burden was accommodated predominantly within established populations, likely through enlargement of pre-existing aggregates rather than continual generation of new aggregates.

One explanation is that glia initially possesses greater capacity to accommodate mHTT through protein quality-control mechanisms. Glia exhibit robust autophagic, lysosomal, phagocytic, and stress-response pathways that handle intracellular and extracellular proteotoxic material ((Abjean et al., 2023; Benraiss et al., 2021; Li & Haney, 2020; Wilson et al., 2023)). Consistently, enhancing glial proteostatic pathways suppresses polyglutamine aggregation and improves glial function ((Yeh et al., 2018)), while glia preferentially generates mature amyloid-β fibrils, whereas neurons accumulate smaller intracellular species associated with greater toxicity (Jonson et al., 2018). Thus, cellular context may influence both protein clearance and aggregate maturation.

This distinct aggregation pattern reveals an apparent disconnect between intracellular mHTT burden and phenotypic severity in glia. Despite progressively increasing HTT immunoreactivity, behavioural impairment remained relatively modest early, contrasting with neuronal mHTT, where rapid accumulation accompanied dynamic aggregate remodelling and early motor dysfunction. Aggregate burden alone may therefore not determine dysfunction; toxicity also depends on the biochemical state, intracellular localization, and structural organization of mHTT species ((Arrasate et al., 2004; Donaldson et al., 2026; Riguet et al., 2021)). Soluble oligomeric and intermediate species are generally considered more biologically reactive than large, highly ordered inclusions, which may represent relatively stable end-stage assemblies. Stable glial aggregate architecture therefore raises the possibility that mHTT is progressively incorporated into existing aggregates, limiting smaller, potentially more toxic intermediates and delaying overt dysfunction. Indeed, huntingtin exon-1 can follow multiple aggregation pathways depending on its molecular environment, while biochemical and subcellular context influences aggregate composition, maturation, and toxicity (Chang et al., 2024; Pandey et al., 2018; Riguet et al., 2021).

However, this interpretation requires caution. The present study does not distinguish soluble from insoluble mHTT, determine aggregate ultrastructure, measure aggregate turnover or proteasomal/autophagic flux, or establish the biochemical properties or toxicity of individual aggregate classes. Aggregate enlargement therefore cannot establish protective sequestration. Alternatively, progressive impairment or saturation of glial degradative pathways could limit mHTT clearance, causing continued accumulation within established aggregates ((Abjean et al., 2023; Al-Dalahmah et al., 2020; Benraiss et al., 2021)). Aggregate retention might also reduce soluble aggregation-prone species available for interaction with neighbouring cells, consistent with glial roles in pathogenic-protein uptake, processing, and compartmentalisation, although this remains untested ((Abjean et al., 2023; Babcock & Ganetzky, 2015; Donnelly et al., 2020)). Thus, the data cannot distinguish protective sequestration from impaired degradation but reveal an important paradox: progressive intracellular mHTT accumulation does not immediately translate into severe behavioural dysfunction, indicating that aggregate burden–toxicity relationships depend strongly on cellular context.

Importantly, stable aggregate architecture does not imply stable glial function. Progressive HTT accumulation, declining *repo* immunoreactivity, and delayed behavioural deterioration indicate continued glial dysfunction despite comparatively stable aggregate organization. Together, these findings support fundamentally different proteostatic trajectories in neurons and glia and raise the next question: how progressive glial pathology contributes to the eventual emergence of severe behavioural dysfunction.

### Progressive neuronal involvement during glial mHTT expression supports non-cell-autonomous disease progression

If glia initially accommodates accumulating mHTT, a key question is how this pathology evolves. A notable finding was the progressive increase in neuronal HTT association following glial mHTT expression. In *repo*>HTTex1pQ93 brains, *elav*–HTT colocalization increased significantly with age, whereas *repo*–HTT colocalization remained relatively unchanged. Thus, despite continued glial mHTT accumulation, an increasing proportion of detectable HTT became associated with neuronal compartments. This shift coincided with increasing intracellular mHTT, declining *repo* immunoreactivity, and delayed severe behavioural deficits, placing progressive neuronal involvement downstream of primary glial pathology.

The neuronal reference showed a distinct pattern: HTT remained preferentially associated with *elav*-positive compartments throughout disease progression. Although repo–HTT overlap varied with age, it broadly paralleled neuronal HTT association, and the absence of a significant age × cell-type interaction indicated no selective shift toward glial compartments. This does not imply complete neuronal confinement, as reactive glial responses and neuron-to-glia mHTT transmission have been reported in HD models ((Abjean et al., 2023; Babcock & Ganetzky, 2015; Donnelly et al., 2020)). Rather, neuronal association remained dominant and its relative distribution between neuronal and glial compartments largely stable within the analysed period.

Importantly, the Manders’ M2 analysis provides the primary quantitative evidence for progressive neuronal HTT association during glia-initiated pathology. The significant age-dependent increase in *elav*–HTT overlap demonstrates increasing spatial association of HTT with neuronal compartments, but cannot distinguish intracellular neuronal mHTT from extracellular, perineuronal, or closely apposed glial signal. Thus, the M2 data establish progressive neuronal involvement, not direct glia-to-neuron transfer. Correlative Holo-CT and fluorescence imaging of a representative *repo*>HTTex1pQ93 brain provided qualitative anatomical support for the M2 findings. Discrete HTT-positive structures were closely associated with *elav*-positive neuronal compartments, with several appearing localized within *elav*-positive cellular domains. Thus, Holo-CT anatomically complements the quantitative M2 analysis by visualizing HTT-positive structures within or closely associated with neuronal domains during glia-initiated pathology. Although restricted to a representative age, this spatial evidence strengthens the interpretation of neuronal HTT association, while establishing temporal redistribution or direct glia-to-neuron transfer would require longitudinal or lineage-tracing approaches.

Nevertheless, several mechanisms could account for this pattern. Intercellular mHTT movement has been reported through extracellular vesicles, tunnelling nanotubes, synaptic transmission, and other forms of cellular communication ((Babcock & Ganetzky, 2015; Donnelly et al., 2020; Victoria & Zurzolo, 2017)). Phagocytic glia can act as obligatory intermediates in mHTT aggregate transmission across neuronal synapses, demonstrating their capacity for pathogenic-protein uptake, processing, and propagation ((Donnelly et al., 2020)). Exopher-mediated disposal may similarly enable stressed cells to release proteotoxic material for uptake and processing by neighbouring cells ((Melentijevic et al., 2017; Trajkovic et al., 2017)). The increasing neuronal HTT association observed here could therefore reflect neuronal uptake of glia-derived mHTT, extracellular HTT redistribution, enlargement of glial aggregates into neuronal territories, or remodelling of neuron–glia interfaces. The present imaging cannot distinguish these possibilities or determine whether associated HTT is intracellular, extracellular, or membrane-associated. Establishing transfer will require lineage-specific protein tracing, photoconvertible reporters, or cell-type-specific biochemical approaches.

Despite these limitations, the temporal pattern is biologically informative. Unlike the neuronal reference, where mHTT remained predominantly neuron-associated, glial expression was accompanied by progressively increasing neuronal HTT association. Thus, glia-initiated pathology increasingly extends beyond the original expressing cell population as disease progresses, consistent with the emergence of a non-cell-autonomous component. This provides a temporal link between progressive glial pathology, increasing neuronal involvement, and the delayed emergence of severe behavioural dysfunction.

### Glial cellular pathology temporally parallels behavioural progression

The behavioural progression closely paralleled the evolving cellular pathology following glial mHTT expression. Although these temporal relationships do not establish causality, they define a coherent sequence linking mHTT accumulation with functional decline. Notably, severe behavioural impairment developed while overall brain architecture remained largely preserved, indicating that cellular dysfunction precedes overt neuroanatomical degeneration.

Glial mHTT followed a gradual, multistage trajectory. Early intracellular accumulation occurred while glial organization and motor performance remained comparatively preserved. This was followed by declining *Repo* immunoreactivity, continued mHTT accumulation within established aggregate populations, increasing neuronal HTT association, and ultimately severe climbing and flight deficits with reduced survival. This progression is consistent with models in which glial dysfunction contributes to neurodegeneration through progressive disruption of neuron–glia interactions ((Benraiss et al., 2021; Villanueva et al., 2025)). Although causality between these events cannot be established here, their temporal correspondence places progressive glial pathology before increasing neuronal involvement and delayed functional deterioration.

The neuronal reference reinforces the distinct trajectory of glia-initiated pathology. Direct neuronal mHTT expression produced rapid intracellular accumulation, dynamic aggregate remodelling, persistent neuronal HTT localization, and early behavioural impairment and mortality, consistent with a predominantly cell-autonomous process ((Creus-Muncunill & Ehrlich, 2019; Donaldson et al., 2026)). In contrast, glial mHTT followed a slower, multistage trajectory characterized by early intracellular accumulation, progressive glial alteration, continued accumulation within established aggregate populations, increasing neuronal HTT association, and ultimately delayed but severe behavioural decline. Thus, the same pathogenic protein follows distinct temporal trajectories depending on its cellular origin. Integrating the behavioural, fluorescence, aggregate, and colocalization analyses, this framework supports glial pathology as an active component of HD-like disease progression rather than merely a secondary response to neuronal pathology.

## Conclusion

The present study demonstrates that glial mHTT expression is sufficient to establish a progressive HD-like pathological trajectory in vivo. Rather than producing a milder version of neuronal pathology, glial HTTex1-Q93 followed a distinct temporal course characterized by progressive intracellular mHTT accumulation within comparatively stable aggregate populations, declining glial homeostasis, increasing neuronal HTT association, and ultimately severe neurological dysfunction. The established neuronal condition, characterized by rapid mHTT accumulation, dynamic aggregate remodelling, and early behavioural decline, provides a reference that highlights the distinct evolution of glia-initiated pathology.

By integrating longitudinal behavioural analyses with quantitative confocal imaging, fluorescence measurements, aggregate architecture, and cell-type-specific colocalization, this study establishes a temporal framework linking glial mHTT accumulation to progressive glial alteration, increasing neuronal involvement, and delayed functional decline. These findings position glial pathology as an active component of HD-like disease progression rather than merely a secondary response to neuronal dysfunction. Future studies defining the molecular basis of glial mHTT handling, increasing neuronal HTT association, and the transition from glial pathology to neuronal involvement will be important for understanding disease progression and identifying cell-type-specific therapeutic targets.

## Materials and Methods

### Drosophila Stocks, Crosses, And Culture Conditions

Transgenic lines w; P{UAS-Httex1p Q20}, w; P{UAS-Httex1p Q93}4F1, pan-neuronal w; P{w+mW.hs = GawB}*elav*C155, and pan-glial *repo*-GAL4/TM6B were used. Human Huntingtin exon-1 was expressed using GAL4/UAS by crossing virgin *elav*-GAL4 or *repo*-GAL4 females with UAS-Httex1p Q20 or Q93 males, generating *elav*>HTTex1pQ20, *elav*>HTTex1pQ93, *repo*>HTTex1pQ20, and *repo*>HTTex1pQ93. Q20 served as the corresponding control for transgene-expression effects independent of polyglutamine expansion. Stocks were obtained from Bloomington Drosophila Stock Center and maintained in the laboratory.

Flies were maintained on standard cornmeal–yeast–agar medium at 25 ± 1°C, 60–70% relative humidity, and a 12 h:12 h light/dark cycle. Parents were transferred to fresh food every 2–3 days to prevent overcrowding and synchronize progeny. Adults were collected within 24 h of eclosion and maintained by genotype at 25–30 flies/vial, with transfer to fresh food every 2–3 days. Only healthy, normally eclosed flies without visible developmental abnormalities were used for behavioural, survival, and imaging experiments.

### Experimental Design

The experimental workflow is summarized in Figure I. Neuronal and glial HTTex1pQ20 and HTTex1pQ93 expression was assessed by behavioural phenotyping and quantitative confocal imaging. Behavioural analyses comprised larval feeding and crawling, eclosion, adult survival, climbing, and flight assays.

Adult brains were analysed at Days 3, 7, 11, 14, and 17 post-eclosion for *repo*>HTTex1pQ93 and controls, and Days 3, 7, 11, and 14 for *elav*>HTTex1pQ93 and controls because of earlier onset and shorter lifespan in the neuronal model. Whole-mount brains were immunostained for *elav*, *repo*, and Huntingtin and imaged by confocal microscopy using identical acquisition settings within each experimental series. FIJI/ImageJ analysis of Z-stacks included corrected mean fluorescence intensity (MFI), three-dimensional aggregate quantification using 3D Object Counter, and thresholded Manders’ colocalization using JACoP.

### Behavioural Assays

Behavioural analyses were performed under identical culture conditions at 25°C using independent biological cohorts generated from separate genetic crosses. Unless otherwise stated, assays were performed as previously described ((Chongtham & Agrawal, 2016)) with minor modifications.

### Larval Crawling Assay

Synchronized wandering third-instar larvae were rinsed in PBS and individually placed on 2% agarose plates over graph paper. Distance travelled in 30 s was determined from grid boxes crossed, with diagonal movements scored as 1.5 boxes. Each larva was tested thrice. Two independent biological replicates of 10 larvae/genotype were analysed, and mean distance per larva was calculated.

### Larval Feeding Assay

Synchronized feeding third-instar larvae were starved in 5% sucrose for 1 h and transferred to 2% (w/v) agarose plates with a central well containing yeast paste supplemented with 4% (w/v) FD&C Blue Dye No. 1 (Sigma-Aldrich). After 2 h feeding, larvae were washed with distilled water and individually homogenized in 200 μL 1× PBS. Homogenates were centrifuged at 13,500 rpm for 10 min and supernatant absorbance measured at 625 nm. Three independent biological replicates of 10 larvae/genotype were analysed, with each larva measured once.

### Adult Eclosion Assay

Eggs collected during a 4-h oviposition period were transferred to fresh food (∼100 eggs/vial; five replicate vials/genotype). Adults were counted daily until eclosion ceased. Eclosion was calculated as:

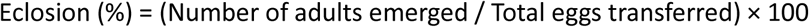

Developmental arrest at different pupal stages was recorded qualitatively (supplementary figure I)

### Survival Assay

Adults collected within 24 h of eclosion were maintained on standard cornmeal medium at 20 flies/vial and transferred to fresh food every 2–3 days. Mortality was recorded daily until all flies died. At least 100 flies/genotype (five replicates of 20) were analysed. Kaplan–Meier survival curves were compared using the log-rank (Mantel–Cox) test.

### Adult Climbing Assay

Negative geotaxis was assessed in groups of 10 flies in transparent vertical tubes (2.2 cm diameter). After 2 min acclimatization, flies were tapped to the bottom and those climbing 10 cm within 15 s were counted. Each biological replicate comprised three technical trials separated by 1 min recovery; three independent biological replicates/genotype/time point were analysed. Performance was expressed as the percentage crossing 10 cm.

### Adult Flight Assay

Flight was assessed using a modified assay described by (Raj, 2015). Groups of 10 flies were starved for 1 h and placed in an empty vial at the centre of a 14.5-cm-diameter beaker half-filled with water, with its inner wall coated with yeast paste and enclosed within a 23-cm-diameter transparent chamber. After 1 h, flies were released. Flies crossing the water and landing on the beaker wall or enclosure were scored as flyers; those remaining in the vial or falling into the water as non-flyers. Three independent biological replicates/genotype/time point were analysed, and flight performance was expressed as percentage flyers.

### Brain Immunohistochemistry

Adult *Drosophila melanogaster* brains were dissected at indicated post-eclosion ages in ice-cold 1× PBS (pH 7.4) using a modified whole-mount protocol ((Kelly et al., 2017)). Anaesthetised flies were immersed in ice-cold PBS, the head capsule opened ventrally beneath the proboscis, and the exoskeleton, retina, trachea, and connective tissues removed while preserving the central brain and optic lobes. Brains were fixed immediately in 4% PFA in 1× PBS for 20 min at room temperature, washed three times for 15 min in 0.2% PBT (PBS, 0.2% Triton X-100), and blocked in 5% BSA/PBT for 3 h at room temperature.

Brains were incubated with primary antibodies in 1× PBS for 12–18 h at 4°C with gentle agitation, washed three times in 0.2% PBT, and incubated with Alexa Fluor-conjugated secondary antibodies in 5% BSA/PBT for 2 h at room temperature in the dark. After three PBT washes, brains were mounted consistently in Vectashield Antifade Mounting Medium containing DAPI (Vector Laboratories), coverslipped, sealed, and stored at 4°C in the dark for ≥12 h before imaging. All genotypes and time points within each staining series were processed simultaneously under identical conditions. Controls lacking primary antibodies assessed non-specific secondary staining. Unless stated otherwise, three biologically independent brains/genotype/time point were analysed.

### Primary and Secondary Antibodies

Triple immunofluorescence was performed to simultaneously visualize neurons, glia, and mutant Huntingtin (mHTT). Primary and secondary antibodies used are listed in Table I. Species-specific, highly cross-adsorbed Alexa Fluor secondary antibodies (Thermo Fisher Scientific) were used to minimize cross-reactivity. Alexa Fluor 488, Alexa Fluor 568, and Alexa Fluor 647 were used to detect *elav*-, *repo*-, and Huntingtin-immunoreactivity, respectively.

**Table I.** Primary and secondary antibodies used for triple immunofluorescence. Primary antibodies against the neuronal marker *elav*, the glial marker *repo*, and mutant Huntingtin (mHTT), together with the corresponding species-specific, highly cross-adsorbed Alexa Fluor-conjugated secondary antibodies, used for whole-mount immunostaining of adult *Drosophila* brains. All antibodies were diluted in blocking solution at the indicated concentrations.

| Antibody | Host species | Supplier | Catalogue no. | Dilution |
| --- | --- | --- | --- | --- |
| Anti-*elav* | Rat | Developmental Studies Hybridoma Bank (DSHB) | Rat-*elav*-7E8A10 (RRID: AB_528218) | 1:200 |
| Anti-*repo* | Mouse | Developmental Studies Hybridoma Bank (DSHB) | 8D12 (RRID: AB_528448) | 1:150 |
| Anti-Huntingtin (mHTT; clone EPR5526) | Rabbit | Novus Biologicals | NBP1-95716 | 1:3000 |
| Donkey anti-rat IgG (H+L), highly cross-adsorbed | Donkey | Thermo Fisher Scientific | A48269 | 1:250 |
| Goat anti-mouse IgG (H+L), highly cross-adsorbed | Goat | Thermo Fisher Scientific | A-11031 | 1:250 |
| Goat anti-rabbit IgG (H+L), highly cross-adsorbed | Goat | Thermo Fisher Scientific | A-21245 | 1:250 |

Alexa Fluor 488, Alexa Fluor 568, and Alexa Fluor 647 were used to detect *elav*-, *repo*-, and Huntingtin-immunoreactivity, respectively.

### Holotomography and Fluorescence Imaging

For correlative holotomography and fluorescence imaging, immunostained whole-mount adult Drosophila brains were transferred to an imaging chamber flushed with 1X PBS, and positioned within the field of view. Imaging was performed using a Tomocube HT-X1™ system (Tomocube Inc., Daejeon, South Korea). Multiple holographic projections acquired at different illumination angles were computationally reconstructed to generate three-dimensional refractive-index tomograms, providing label-free structural information. Fluorescence images of *elav*, *repo*, and HTT were acquired and spatially overlaid with the reconstructed holotomographic data. For visualization, an optimal whole-brain Z-plane was selected, and sequential Z-planes from defined regions of interest were examined to assess the three-dimensional spatial association of HTT-positive structures with neuronal and glial compartments. Holo-CT imaging was performed on a representative Day 14 *repo*>HTTex1pQ93 brain and was used as qualitative spatial validation rather than for quantitative or longitudinal analysis.

### Confocal Microscopy

Immunostained adult brains were imaged on a Leica TCS SP8 confocal microscope (Leica Microsystems, Germany) with white light laser (WLL), hybrid detectors, and LAS X v3.5.7, using an HC PL APO CS2 20×/0.75 NA dry objective and identical settings within each staining batch. Alexa Fluor 488 (*elav*), 568 (*repo*), and 647 (HTT) were sequentially imaged at 500–550, 570–625, and 650–700 nm, respectively. Whole-brain Z-stacks were acquired at 1024 × 1024 pixels (0.28 μm/pixel), with Z-step intervals optimized by LAS X.

XYZ images were acquired unidirectionally at 400 Hz, 1 Airy unit, 1× digital zoom, and 0.7875 μs pixel dwell time without averaging. Alexa Fluor 488 was excited at 488 nm (2% WLL power); Alexa Fluor 568 and 647 used the corresponding WLL lines. AOBS-controlled spectral detection minimized crosstalk. Laser power, detector gain/offset, pinhole, scan speed, zoom, pixel size, and spectral settings were constant within each experimental series. Quantitative fluorescence and aggregate analyses were restricted to within-paradigm comparisons; neuronal and glial models were compared qualitatively.

### Image Processing and Quantitative Analysis

Raw stacks were exported without deconvolution or intensity enhancement and analysed in FIJI/ImageJ using established workflows ((Brazill et al., 2018; Roth et al., 2023)). Identical whole-brain ROIs were manually defined. Cell-free background fluorescence was subtracted from channel MFI for each optical section, and corrected MFI was averaged across the Z-stack to yield one value/brain. Maximum-intensity projections were used only for representative figures.

HTT colocalization with *elav* or *repo* was analysed independently using JACoP. Manually defined Costes thresholds, applied consistently within each experimental group, were used to calculate thresholded Manders’ coefficients, with M2 as the primary measure of HTT association with neuronal or glial compartments. Costes randomization (1000 iterations) assessed colocalization above random expectation ((BOLTE & CORDELIÈRES, 2006)).

HTT aggregates were quantified using 3D Object Counter after intensity thresholding and three-dimensional segmentation (Bolte & Cordelières, 2006; Brazill et al., 2018). Three-dimensional voxel connectivity identified individual aggregates, and object number, centroid coordinates, volume, voxel count, surface area, and integrated fluorescence intensity were exported per brain.

Aggregate volumes from all experimental brains were pooled to establish fixed percentile-derived thresholds applied uniformly to all samples. Aggregates were classified as puncta (<10 μm³), small (10– 32 μm³), large (32–132 μm³), or inclusions (>132 μm³). Aggregate number, cumulative volume, and composition were calculated per class using identical processing, segmentation, and classification parameters within each experimental series. The workflow is summarized in Figure II.

### Quality Control

Identical image-processing, segmentation, and analysis parameters were used within each experimental series. Aggregate thresholds were defined from the pooled global volume distribution and applied uniformly. Quantification used complete three-dimensional stacks; maximum-intensity projections were used only for representative figures.

### Statistical Analysis

Analyses were performed in GraphPad Prism 10 (GraphPad Software, San Diego, CA, USA). Data are mean ± SEM, with independent fly cohorts or individual brains as biological replicates. P < 0.05 was considered significant.

Larval crawling and eclosion were analysed by ordinary one-way ANOVA with Tukey’s multiple-comparisons test; feeding by unpaired two-tailed Student’s *t*-test; climbing and flight by ordinary two-way ANOVA (genotype × age) with Tukey’s test; and survival by Kaplan–Meier analysis with the log-rank (Mantel–Cox) test.

Corrected *elav*, *repo*, and HTT MFI was analysed separately for each expression paradigm by ordinary two-way ANOVA (genotype × age) with Šídák’s multiple-comparisons test. Thresholded Manders’ M2 was analysed by ordinary two-way ANOVA (cellular compartment × age) with Tukey’s test where appropriate; Costes randomization (1000 iterations) validated colocalization above random overlap. Aggregate number and cumulative volume were analysed separately for each expression paradigm by ordinary two-way ANOVA (aggregate class × age) with Tukey’s multiple-comparisons test.

## Author Contributions

**Debalina Chatterjee:** Conceptualization, Methodology, Investigation, Formal analysis, Data curation, Visualization, Writing – original draft, review & editing. **Jassika Gupta**: Data Visualization, Writing – review, **Mansi Yadav**: Data Visualization, **Parul Mittal**: Writing – review, **Namita Agrawal:** Supervision, Resources, Writing – review & editing. **Anju Shrivastava:** Conceptualization, Supervision, Writing – review & editing.

## Compliance with Ethical Standards

This study used *Drosophila melanogaster* and did not involve human participants or vertebrate animals. Institutional ethical approval was therefore not required.

## Data Availability

The datasets generated and analysed during the current study are available from the corresponding author upon reasonable request.

## AI Use Disclosure

During the preparation of this manuscript, the authors used ChatGPT (OpenAI) and Grammarly to assist with language refinement, grammar, and manuscript organization. All scientific content, data interpretation, analyses, and conclusions were developed and verified by the authors. The authors critically reviewed, revised, and approved the final manuscript and accept full responsibility for its content.

---

**Figure.**
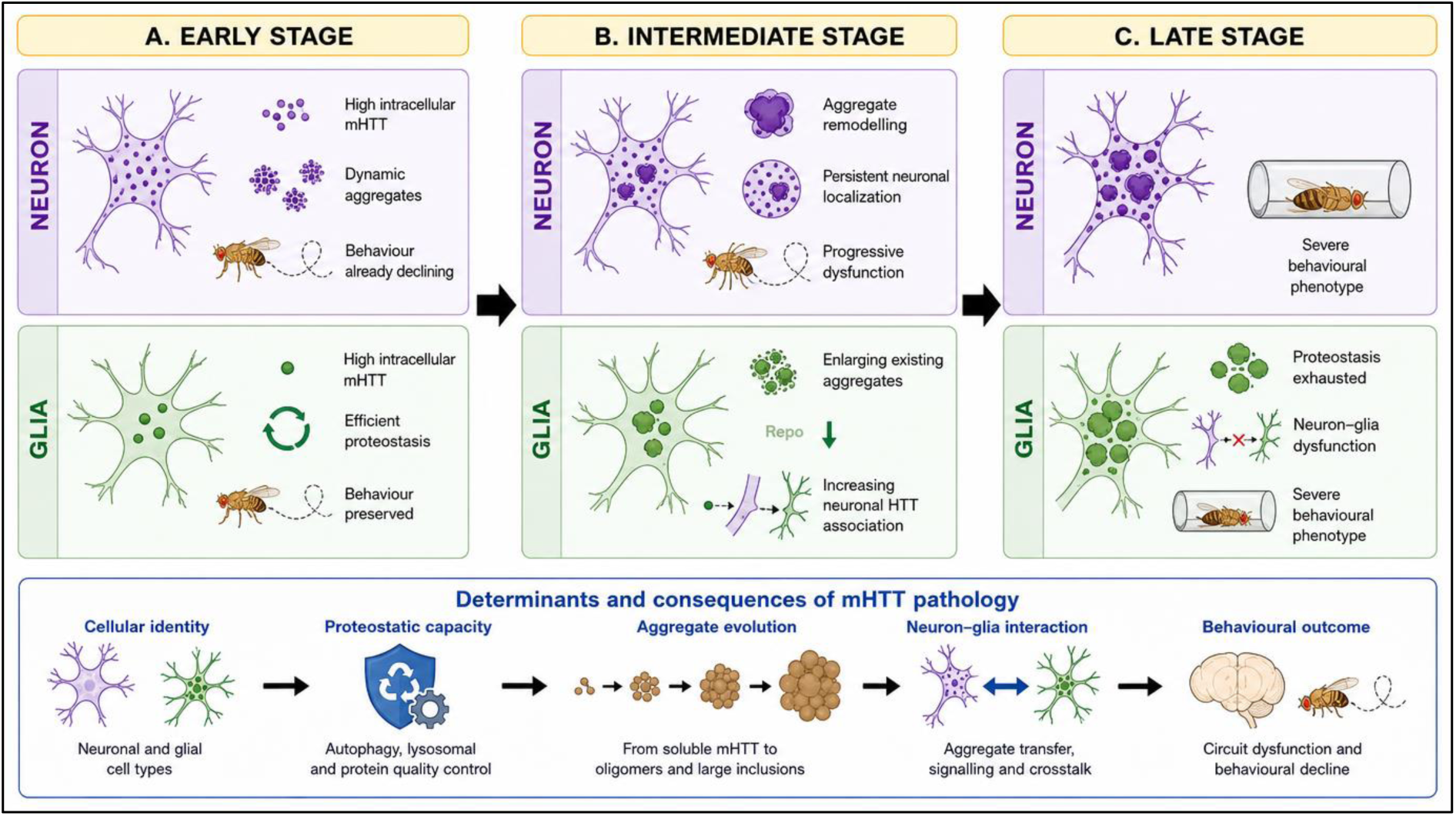
Proposed model illustrating the cell-type-specific temporal trajectories of mutant Huntingtin pathology in Drosophila. The schematic summarizes the principal findings of the present study and proposes a model in which the cellular context of mutant Huntingtin (mHTT) expression determines the temporal progression of Huntington’s disease pathology. (A) Early stage: Pan-neuronal expression of mHTT (*elav*>HTTex1pQ93) results in rapid intracellular accumulation of the mutant protein, dynamic aggregate formation, and early behavioural impairment, consistent with a predominantly cell-autonomous pathological process. In contrast, pan-glial expression (*repo*>HTTex1pQ93) also leads to intracellular mHTT accumulation but is initially accompanied by preserved behavioural performance, suggesting effective proteostatic buffering by glial cells. (B) Intermediate stage: Neuronal pathology is characterized by continuous aggregate remodelling, persistent neuronal localization of mHTT, and progressive functional decline. In the glial model, existing aggregates enlarge progressively while *Repo* immunoreactivity declines, indicating deterioration of glial integrity. Concurrently, increasing neuronal association of mHTT suggests the emergence of non-cell-autonomous interactions between glia and neurons. (C) Late stage: Neuronal expression culminates in severe behavioural deficits associated with sustained intracellular mHTT pathology. In the glial model, progressive exhaustion of glial proteostatic capacity and disruption of neuron–glia homeostasis are proposed to ultimately converge on behavioural impairment comparable to that observed following neuronal expression. The lower panel summarizes the conceptual framework emerging from this study, whereby cellular identity influences proteostatic capacity, which in turn governs aggregate evolution, neuron–glia interactions, and ultimately behavioural outcome. This model proposes that identical pathogenic mHTT follows distinct temporal trajectories depending on the cell type in which it is expressed, giving rise to either a predominantly cell-autonomous neuronal pathway or a delayed, multistage glial-mediated route to neurodegeneration.

**Figure I.**
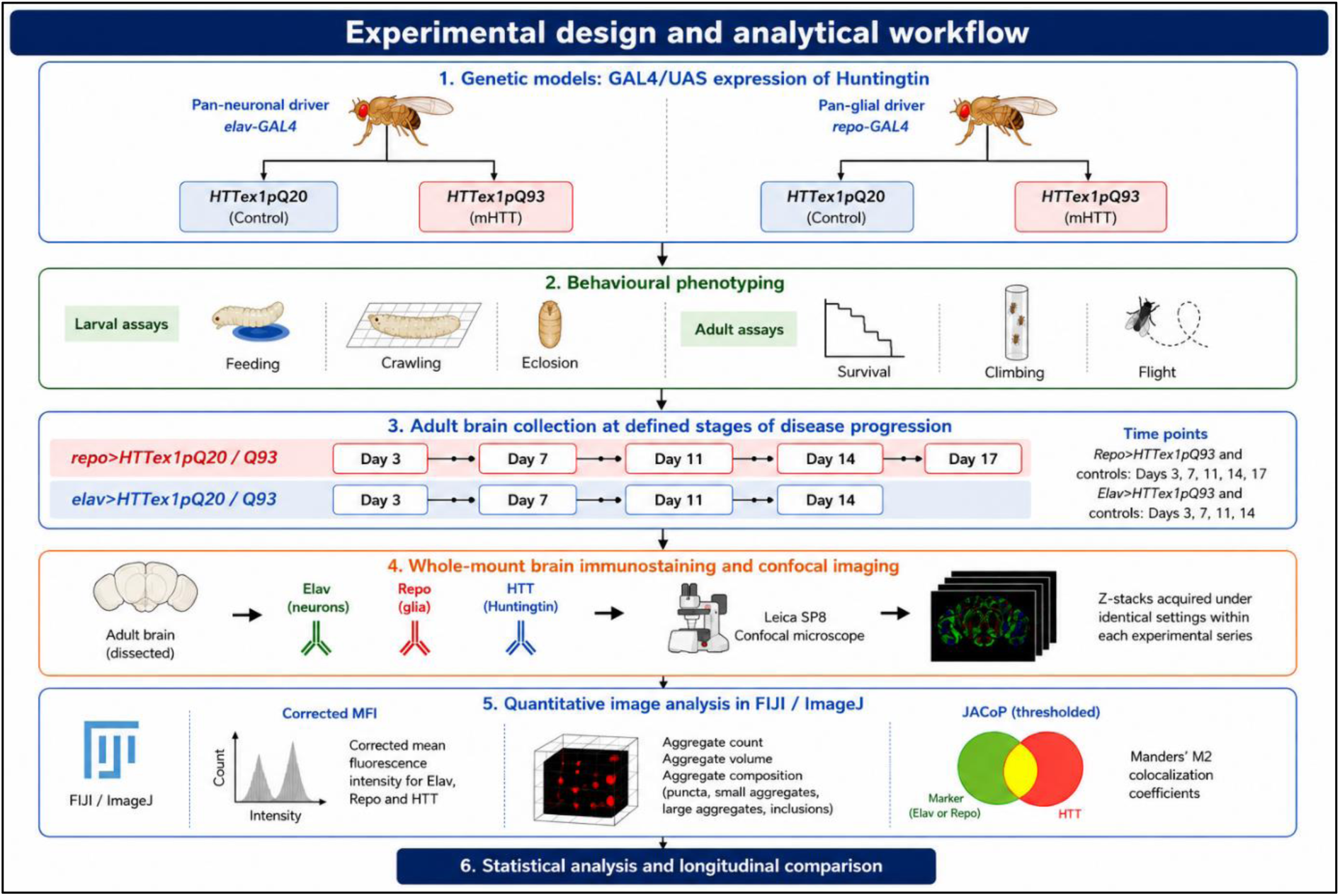
Experimental design and analytical workflow. Adult *Drosophila melanogaster* models were generated using the GAL4/UAS system to express wild-type Huntingtin exon-1 (HTTex1pQ20) or mutant Huntingtin exon-1 (HTTex1pQ93) pan-neuronally (*elav*-GAL4) or pan-glially (*repo*-GAL4). Disease progression was assessed by combining behavioural phenotyping with longitudinal quantitative confocal imaging. Behavioural analyses included larval feeding, larval crawling, eclosion, adult survival, climbing, and flight assays. Adult brains were collected at defined post-eclosion time points corresponding to disease progression (*repo*>HTTex1pQ20/Q93: Days 3, 7, 11, 14, and 17; *elav*>HTTex1pQ20/Q93: Days 3, 7, 11, and 14). Whole-mount brains were immunostained for the neuronal marker *elav*, the glial marker *Repo*, and mutant Huntingtin (mHTT), followed by confocal microscopy using identical acquisition settings within each experimental series. Quantitative image analysis of confocal Z-stacks in FIJI/ImageJ included corrected mean fluorescence intensity (MFI), three-dimensional aggregate quantification using the 3D Object Counter plugin, and cell-type-specific colocalization analysis using JACoP, with thresholded Manders’ M2 coefficient as the primary measure of HTT association. The integrated behavioural and imaging datasets were subsequently analysed to define the temporal progression of neuronal-and glial-driven Huntington’s disease pathology.

**Figure II.**
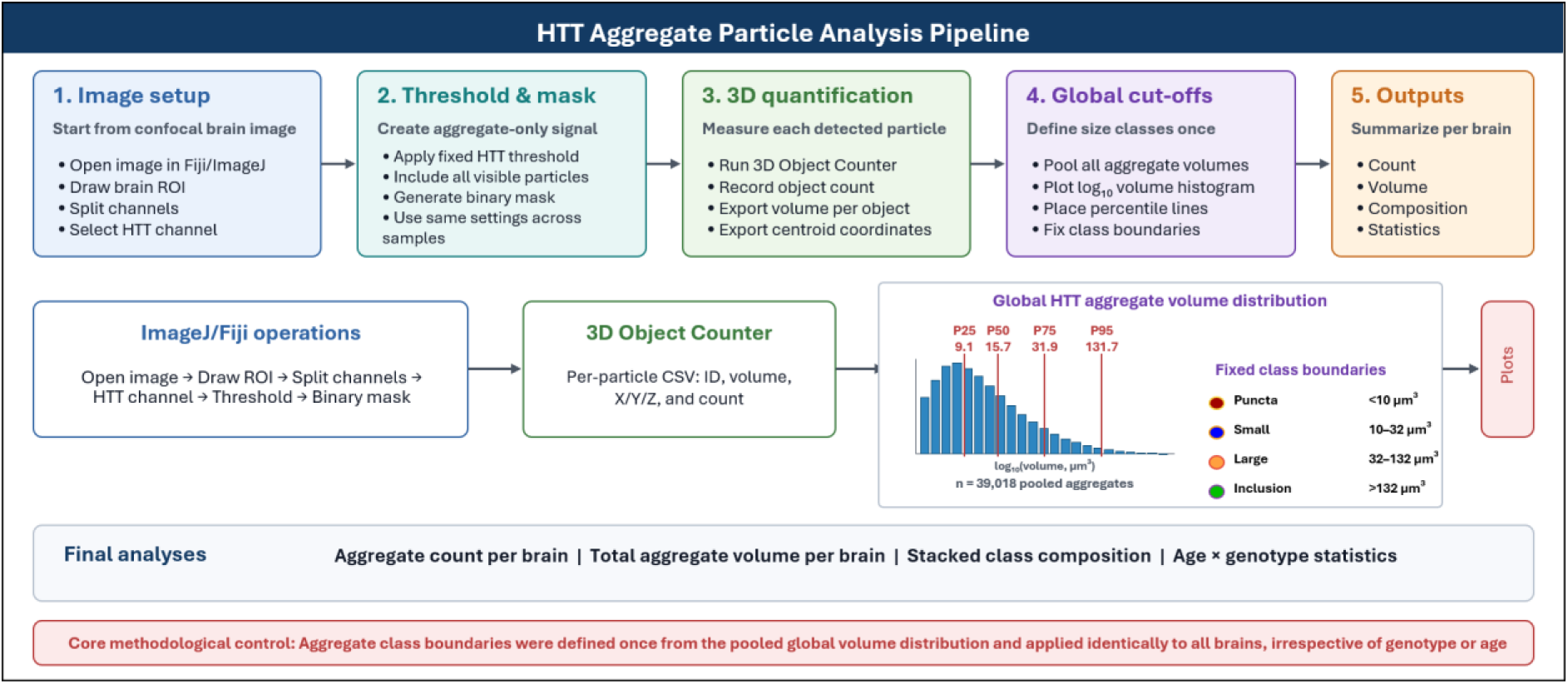
Workflow for quantitative three-dimensional analysis and classification of mutant huntingtin aggregates. Schematic representation of the image analysis pipeline used for quantitative assessment of mutant huntingtin (HTT) aggregates in adult *Drosophila* brains. Confocal Z-stack images were first imported into Fiji/ImageJ, and the brain region of interest (ROI) was selected. Individual fluorescence channels were separated, and the HTT channel was processed using a fixed intensity threshold applied uniformly across all samples to generate binary masks containing HTT-positive aggregates. Three-dimensional aggregate quantification was performed using the 3D Object Counter plugin, which generated per-particle measurements including object number, volume, centroid coordinates, and spatial position. Aggregate volumes from all experimental groups and time points were subsequently pooled to generate a global volume distribution. Percentile-based cut-offs derived from this pooled distribution (25th, 50th, 75th, and 95th percentiles) were used to define four aggregate classes: puncta (<10 μm³), small aggregates (10–32 μm³), large aggregates (32–132 μm³), and inclusions (>132 μm³). These fixed class boundaries were applied uniformly to every brain, independent of genotype or age, to ensure unbiased comparison of aggregate composition across experimental groups. Final quantitative outputs included total aggregate count, cumulative aggregate volume, aggregate class composition, and statistical comparisons across age and genotype. The use of a single global volume distribution to establish aggregate class boundaries ensured that all samples were analysed using identical classification criteria.

## References

1. Abjean, L., Ben Haim, L., Riquelme-Perez, M., Gipchtein, P., Derbois, C., Palomares, M.-A., Petit, F., Hérard, A.-S., Gaillard, M.-C., Guillermier, M., Gaudin-Guérif, M., Aurégan, G., Sagar, N., Héry, C., Dufour, N., Robil, N., Kabani, M., Melki, R., De la Grange, P., … Escartin, C. (2023). Reactive astrocytes promote proteostasis in Huntington’s disease through the JAK2-STAT3 pathway. Brain, 146(1), 149–166. 10.1093/brain/awac068

2. Al-Dalahmah, O., Sosunov, A. A., Shaik, A., Ofori, K., Liu, Y., Vonsattel, J. P., Adorjan, I., Menon, V., & Goldman, J. E. (2020). Single-nucleus RNA-seq identifies Huntington disease astrocyte states. Acta Neuropathologica Communications, 8(1), 19. 10.1186/s40478-020-0880-6

3. Arrasate, M., Mitra, S., Schweitzer, E. S., Segal, M. R., & Finkbeiner, S. (2004). Inclusion body formation reduces levels of mutant huntingtin and the risk of neuronal death. Nature, 431(7010), 805–810. 10.1038/nature02998

4. Babcock, D. T., & Ganetzky, B. (2015). Transcellular spreading of huntingtin aggregates in the Drosophila brain. Proceedings of the National Academy of Sciences of the United States of America, 112(39), E5427–33. 10.1073/pnas.1516217112

5. Bates, G. P. (2025). Somatic CAG-repeat expansion drives neuronal loss in Huntington’s disease. Neuron, 113(3), 342–344. 10.1016/j.neuron.2025.01.008

6. Benraiss, A., Mariani, J. N., Osipovitch, M., Cornwell, A., Windrem, M. S., Villanueva, C. B., Chandler-Militello, D., & Goldman, S. A. (2021). Cell-intrinsic glial pathology is conserved across human and murine models of Huntington’s disease. Cell Reports, 36(1), 109308. 10.1016/j.celrep.2021.109308

7. Bhat, A. A., Moglad, E., Afzal, M., Thapa, R., Almalki, W. H., Kazmi, I., Alzarea, S. I., Ali, H., Pant, K., Singh, T. G., Dureja, H., Singh, S. K., Dua, K., Gupta, G., & Subramaniyan, V. (2024). Therapeutic approaches targeting aging and cellular senescence in Huntington’s disease. CNS Neuroscience & Therapeutics, 30(10). 10.1111/cns.70053

8. Bolte, S., & CordeliÈees, F. P. (2006). A guided tour into subcellular colocalization analysis in light microscopy. Journal of Microscopy, 224(3), 213–232. 10.1111/j.1365-2818.2006.01706.x

9. Brazill, J. M., Zhu, Y., Li, C., & Zhai, R. G. (2018). Quantitative Cell Biology of Neurodegeneration in *Drosophila* Through Unbiased Analysis of Fluorescently Tagged Proteins Using ImageJ. Journal of Visualized Experiments, (138). 10.3791/58041

10. Chang, C.-P., Wu, C.-W., & Chern, Y. (2024). Metabolic dysregulation in Huntington’s disease: Neuronal and glial perspectives. Neurobiology of Disease, 201, 106672. 10.1016/j.nbd.2024.106672

11. Chongtham, A., & Agrawal, N. (2016). Curcumin modulates cell death and is protective in Huntington’s disease model. Scientific Reports, 6(1), 18736. 10.1038/srep18736

12. Creus-Muncunill, J., & Ehrlich, M. E. (2019). Cell-Autonomous and Non-cell-Autonomous Pathogenic Mechanisms in Huntington’s Disease: Insights from In Vitro and In Vivo Models. Neurotherapeutics, 16(4), 957–978. 10.1007/s13311-019-00782-9

13. Cvetanovic, M., & Gray, M. (2023). Contribution of Glial Cells to Polyglutamine Diseases: Observations from Patients and Mouse Models. Neurotherapeutics, 20(1), 48–66. 10.1007/s13311-023-01357-5

14. Donaldson, J., Hensman Moss, D., Ciosi, M., Usdin, K., Balmus, G., Monckton, D. G., & Tabrizi, S. J. (2026). Huntington disease: somatic expansion, pathobiology and therapeutics. Nature Reviews Neurology, 22(1), 5–21. 10.1038/s41582-025-01159-7

15. Donnelly, K. M., DeLorenzo, O. R., Zaya, A. DA, Pisano, G. E., Thu, W. M., Luo, L., Kopito, R. R., & Panning Pearce, M. M. (2020). Phagocytic glia are obligatory intermediates in transmission of mutant huntingtin aggregates across neuronal synapses. ELife, 9. 10.7554/eLife.58499

16. Gao, C., Jiang, J., Tan, Y., & Chen, S. (2023). Microglia in neurodegenerative diseases: mechanism and potential therapeutic targets. Signal Transduction and Targeted Therapy, 8(1), 359. 10.1038/s41392-023-01588-0

17. Goldman, S. A. (2024). The role of glial pathology in Huntington’s disease. In Huntington’s Disease (pp. 337–351). Elsevier. 10.1016/B978-0-323-95672-7.00020-0

18. Jansen, A. H. P., van Hal, M., op den Kelder, I. C., Meier, R. T., de Ruiter, A., Schut, M. H., Smith, D. L., Grit, C., Brouwer, N., Kamphuis, W., Boddeke, H. W. G. M., den Dunnen, W. F. A., van Roon, W. M. C., Bates, G. P., Hol, E. M., & Reits, E. A. (2017). Frequency of nuclear mutant huntingtin inclusion formation in neurons and glia is cell-type-specific. Glia, 65(1), 50–61. 10.1002/glia.23050

19. Jonson, M., Nyström, S., Sandberg, A., Carlback, M., Michno, W., Hanrieder, J., Starkenberg, A., Nilsson, K. P. R., Thor, S., & Hammarström, P. (2018). Aggregated Aβ1-42 Is Selectively Toxic for Neurons, Whereas Glial Cells Produce Mature Fibrils with Low Toxicity in Drosophila. Cell Chemical Biology, 25(5), 595–610.e5. 10.1016/j.chembiol.2018.03.006

20. Kelly, S. M., Elchert, A., & Kahl, M. (2017). Dissection and Immunofluorescent Staining of Mushroom Body and Photoreceptor Neurons in Adult *Drosophila melanogaster* Brains. Journal of Visualized Experiments, (129). 10.3791/56174

21. Li, Q., & Haney, M. S. (2020). The role of glia in protein aggregation. Neurobiology of Disease, 143, 105015. 10.1016/j.nbd.2020.105015

22. Melentijevic, I., Toth, M. L., Arnold, M. L., Guasp, R. J., Harinath, G., Nguyen, K. C., Taub, D., Parker, J. A., Neri, C., Gabel, C. V., Hall, D. H., & Driscoll, M. (2017). C. elegans neurons jettison protein aggregates and mitochondria under neurotoxic stress. Nature, 542(7641), 367–371. 10.1038/nature21362

23. Ojalvo-Pacheco, J., Yakhine-Diop, S. M. S., Fuentes, J. M., Paredes-Barquero, M., & Niso-Santano, M. (2024). Role of TFEB in Huntington’s Disease. Biology, 13(4), 238. 10.3390/biology13040238

24. Olsen, A. L., & Feany, M. B. (2019). Glial α-synuclein promotes neurodegeneration characterized by a distinct transcriptional program in vivo. Glia, 67(10), 1933–1957. 10.1002/glia.23671

25. Pandey, N. K., Isas, J. M., Rawat, A., Lee, R. V., Langen, J., Pandey, P., & Langen, R. (2018). The 17-residue-long N terminus in huntingtin controls stepwise aggregation in solution and on membranes via different mechanisms. Journal of Biological Chemistry, 293(7), 2597–2605. 10.1074/jbc.M117.813667

26. Raj, A.; A. U.; D. H.; L. S. C. (2015). A simple method to assay flight ability in Drosophila. Drosophila Information Service, 98, 18–22.

27. Rangel-Gomez, M., Alberini, C. M., Deneen, B., Drummond, G. T., Manninen, T., Sur, M., & Vicentic, A. (2024). Neuron–Glial Interactions: Implications for Plasticity, Behavior, and Cognition. The Journal of Neuroscience, 44(40), e1231242024. 10.1523/JNEUROSCI.1231-24.2024

28. Riguet, N., Mahul-Mellier, A.-L., Maharjan, N., Burtscher, J., Croisier, M., Knott, G., Hastings, J., Patin, A., Reiterer, V., Farhan, H., Nasarov, S., & Lashuel, H. A. (2021). Nuclear and cytoplasmic huntingtin inclusions exhibit distinct biochemical composition, interactome and ultrastructural properties. Nature Communications, 12(1), 6579. 10.1038/s41467-021-26684-z

29. Roth, J. R., Moraes, R. C. M. de, Xu, B. P., Crawley, S. R., Khan, M. A., & Melkani, G. C. (2023). Rapamycin reduces neuronal mutant huntingtin aggregation and ameliorates locomotor performance in Drosophila. Frontiers in Aging Neuroscience, 15. 10.3389/fnagi.2023.1223911

30. Scarpelli, E. M., Trinh, V. Y., Tashnim, Z., Krans, J. L., Keller, L. C., & Colodner, K. J. (2019). Developmental expression of human tau in Drosophila melanogaster glial cells induces motor deficits and disrupts maintenance of PNS axonal integrity, without affecting synapse formation. PLOS ONE, 14(12), e0226380. 10.1371/journal.pone.0226380

31. Tarasiuk, O., & Scuteri, A. (2022). Role of Tunneling Nanotubes in the Nervous System. International Journal of Molecular Sciences, 23(20), 12545. 10.3390/ijms232012545

32. Tong, H., Yang, T., Xu, S., Li, X., Liu, L., Zhou, G., Yang, S., Yin, S., Li, X.-J., & Li, S. (2024). Huntington’s Disease: Complex Pathogenesis and Therapeutic Strategies. International Journal of Molecular Sciences, 25(7), 3845. 10.3390/ijms25073845

33. Trajkovic, K., Jeong, H., & Krainc, D. (2017). Mutant Huntingtin Is Secreted via a Late Endosomal/Lysosomal Unconventional Secretory Pathway. The Journal of Neuroscience, 37(37), 9000–9012. 10.1523/JNEUROSCI.0118-17.2017

34. Victoria, G. S., & Zurzolo, C. (2017). The spread of prion-like proteins by lysosomes and tunneling nanotubes: Implications for neurodegenerative diseases. Journal of Cell Biology, 216(9), 2633– 2644. 10.1083/jcb.201701047

35. Villanueva, C. B., Huynh, N. P. T., Mariani, J. N., Mansky, B., Tate, A., Lorenzen, S. S., Chandler-Militello, D., Benraiss, A., & Goldman, S. A. (2025). Human glial progenitors transplanted into Huntington disease mice normalize neuronal gene expression, dendritic structure, and behavior. Cell Reports, 44(6), 115762. 10.1016/j.celrep.2025.115762

36. Wen, W., & Turrigiano, G. G. (2024). Keeping Your Brain in Balance: Homeostatic Regulation of Network Function. Annual Review of Neuroscience, 47(1), 41–61. 10.1146/annurev-neuro-092523-110001

37. Wilson, M. R., Satapathy, S., & Vendruscolo, M. (2023). Extracellular protein homeostasis in neurodegenerative diseases. Nature Reviews Neurology. 10.1038/s41582-023-00786-2

38. Wilton, D. K., & Stevens, B. (2020). The contribution of glial cells to Huntington’s disease pathogenesis. Neurobiology of Disease, 143, 104963. 10.1016/j.nbd.2020.104963

39. Yeh, P.-A., Liu, Y.-H., Chu, W.-C., Liu, J.-Y., & Sun, Y. H. (2018). Glial expression of disease-associated poly-glutamine proteins impairs the blood–brain barrier in Drosophila. Human Molecular Genetics, 27(14), 2546–2562. 10.1093/hmg/ddy160

