## Supplementary Figure I for "Glial Mutant Huntingtin Initiates a Progressive Pathological Cascade Toward Neuronal Involvement in Drosophila"

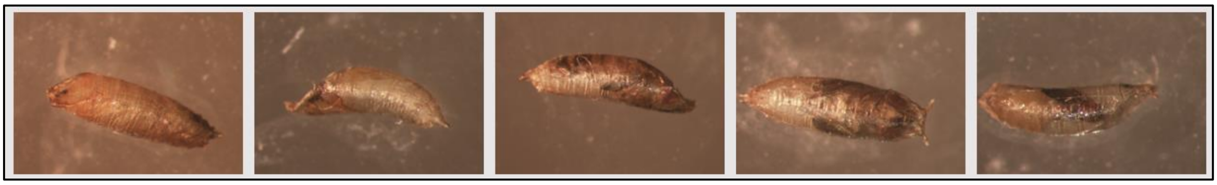

**Supplementary Figure I. Developmental arrest at different pupal stages following glial mHTT expression.** Representative images showing variable stages of pupal developmental arrest observed in *repo*>HTTex1pQ93 animals that failed to complete eclosion. Images qualitatively illustrate the range of developmental abnormalities and were not subjected to quantitative analysis.
